# Characterization of a triad of genes in cyanophage S-2L sufficient to replace adenine by 2-aminoadenine in bacterial DNA

**DOI:** 10.1101/2021.04.30.442174

**Authors:** D. Czernecki, F. Bonhomme, P.A. Kaminski, M. Delarue

**Affiliations:** Unit of Architecture and Dynamics of Biological Macromolecules, CNRS UMR 3528, 25-28 rue du Docteur Roux, Institut Pasteur, 75015 Paris, France; Sorbonne Université, Collège Doctoral, ED 515, 75005 Paris, France; Unit of Epigenetic Chemistry, CNRS UMR 3523, Institut Pasteur, 75015 Paris, France; Unit of Biology of Pathogenic Gram-Positive Bacteria, 25-28 rue du Docteur Roux, Institut Pasteur, 75015 Paris, France

## Abstract

Cyanophage S-2L is known to profoundly alter the biophysical properties of its DNA by replacing all adenines (A) with 2-aminoadenines (Z), which still pair with thymines but with a triple hydrogen bond. It was recently demonstrated that a homologue of adenylosuccinate synthase (PurZ) and a dATP triphosphohydrolase (DatZ) are two important pieces of the metabolism of 2-aminoadenine, participating in the synthesis of ZTGC-DNA. Here, we determine that S-2L PurZ can use either dATP or ATP as a source of energy, thereby also depleting the pool of nucleotides in dATP. Furthermore, we identify a conserved gene (*mazZ)* located between *purZ* and *datZ* genes in *Siphoviridae* phage genomes, and show that it encodes a (d)GTP-specific diphosphohydrolase, thereby providing the substrate of PurZ in the 2-aminoadenine synthesis pathway. High-resolution crystal structures of S-2L PurZ and MazZ with their respective substrates provide a rationale for their specificities. The Z-cluster made of these three genes – *datZ*, *mazZ* and *purZ* – was expressed in *E. coli*, resulting in a successful incorporation of 2-aminoadenine in the bacterial chromosomal and plasmidic DNA. This work opens the possibility to study synthetic organisms containing ZTGC-DNA.

## INTRODUCTION

At least since the last universal common ancestor (LUCA), four types of nucleobases – adenine (A), guanine (G), cytosine (C) and thymine (T) – are used to encode genetic information in the DNA of all living cells (1). This can be extended to viruses, which are important agents of evolution (2). However, the genetic material may be subject to nucleobase modifications, for instance to confer an additional, epigenetic information or to provide resistance against restriction-modification systems. Phages T2, T4 and T6 systematically substitute 5-hydroxymethylcytosine (5hmC) for cytosine (3) protecting the DNA from CRISPR-Cas9 degradation (4), whereas phage 9g replaces a quarter of its genomic guanine with archaeosine (G+) (5), enabling its DNA to resist 71% of cellular restriction enzymes (6).

Such alterations are made almost exclusively without changing the Watson-Crick base-pairing scheme. The only known exception to this rule was revealed by the discovery of cyanophage S-2L, from *Siphoviridae* family (7). S-2L abandons the usage of genomic adenine in favor of 2-aminoadenine (2,6-diaminopurine or Z) (Fig. 1A). The resulting ZTGC-DNA has an improved thermal stability (8, 9) and proves to be almost completely resistant to adenine-targeting restriction enzymes (10).

**Figure 1.**
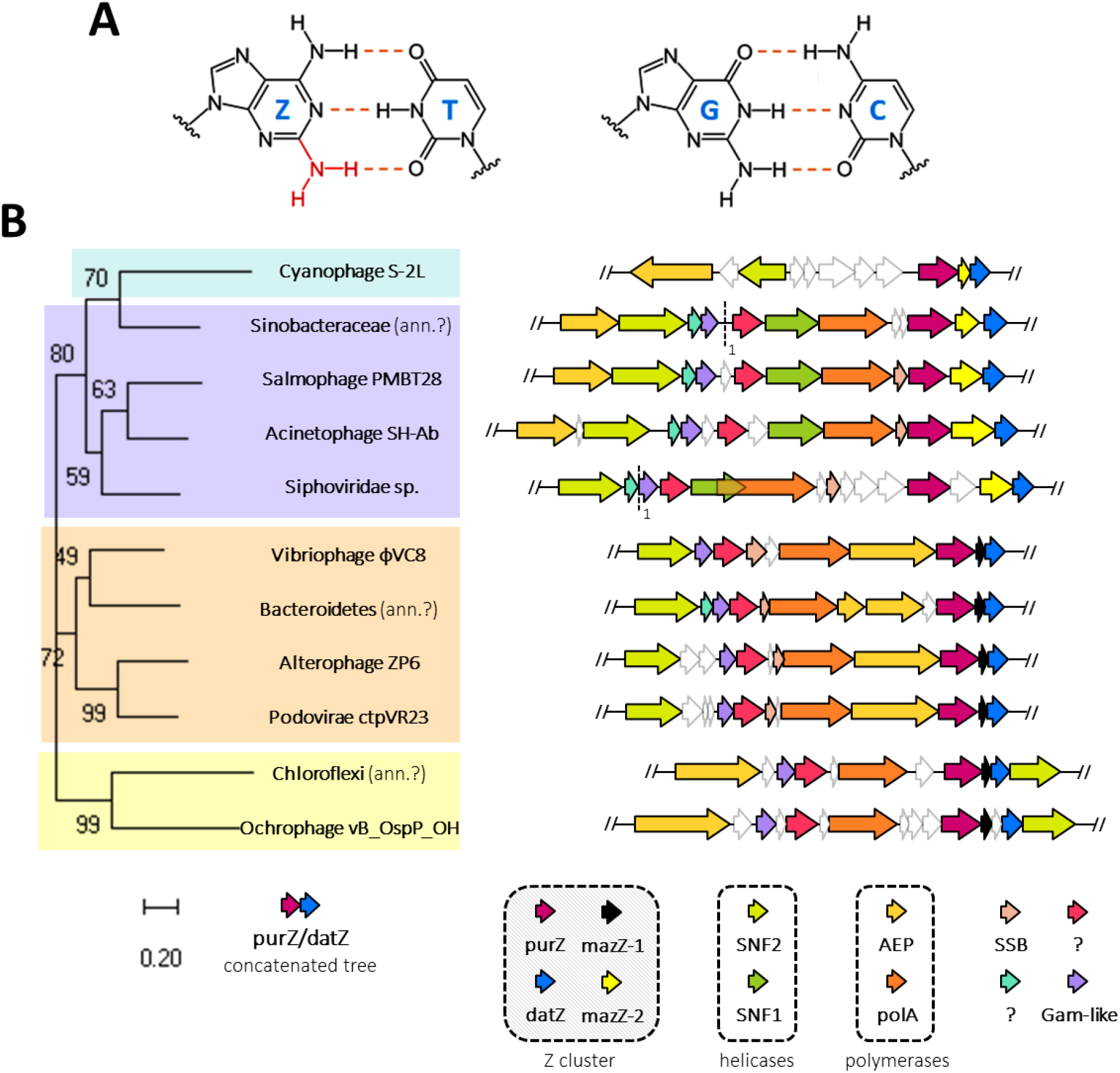
ZTGC-DNA and conservation of the Z-cluster in *Siphoviridae*. **A.**Chemical structure of Z:T and G:C base pairs constituting S-2L’s ZTGC-DNA. **B.** Genomic map of most important replication genes in all phages with close *purZ* and *datZ* homologues, and their phylogeny. The phages can be presently divided into four clades (colours in background to the left) with distinct organisation and variants of replication-related genes. The names of these genes, identified by their colours, are shown below. Cyanophage S-2L is, until now, the only representative of its clade. Close scrutiny of the sequence data reveals possible mis-annotation of some viral sequences with bacterial names (ann. ? note) – their length and content perfectly match other viral sequences and they are probably integrated prophages.

The key enzyme of Z metabolism was recently identified as PurZ, a homologue of adenylosuccinate synthases (PurA) (Sleiman *et al*., in press). PurZ of cyanophage S-2L and related vibriophage ϕVC8 create the immediate precursor of 2-aminoadenine deoxyribose monophosphate (dZMP), N6-succino-2-amino-2’-deoxyadenylate monophosphate (dSMP), from standard dGMP and free aspartic acid (Asp) using ATP as the energy donor. The enzymes necessary for the subsequent conversion of dSMP to dZMP and then dZTP have not been found on the phages’ genomes; instead, the non-specific enzymes of *V. cholerae* complement the pathway of dZTP metabolism of ϕVC8 *in vitro* (Sleiman *et al*., in press).

Furthermore, in S-2L a dATP-specific triphosphatase (DatZ) eliminates dATP from the pool of available dNTPs substrates (11). This step is essential for conferring an indirect nucleotide specificity to the DNA primase-polymerase (PrimPol) of the cyanophage. Co-conservation of both *purZ* and *datZ* genes in related viruses suggest that dATP depletion is closely tied to dZTP production.

Here, we broaden the substrate spectrum of S-2L’s PurZ and present its structure in a complex with dGMP and dATP as an alternative energy donor. Moreover, along with *datZ* and *purZ* we identify a co-conserved gene encoding a MazG-like nucleotide phosphohydrolase (*mazZ*). We show that MazZ generates (d)GMP from (d)GTP, placing the enzyme directly upstream of PurZ in the 2-aminoadenine synthesis pathway. The crystal structure of MazZ bound to the intermediate product allows to identify crucial catalytic residues, shared with close homologues of the enzyme. To confirm the necessity and sufficiency of the Z-cluster (*datZ*, *mazZ*, *purZ*) for efficient ZTGC-DNA synthesis *in cellulo*, we expressed these three genes in *E. coli*. Although toxic to the bacteria, the system was able to convert a significant amount of DNA’s adenine to 2-aminoadenine both in chromosomal and plasmidic fractions.

## RESULTS

### Genome of cyanophage S-2L and its relatives: a conserved Z-cluster

Although the genome of phage S-2L was made available in 2004 (GenBank AX955019), homology analysis suggested several sequence errors. We decided to re-sequence it using the next-generation sequencing (NGS) technology (GenBank MW334946) with improved quality (Suppl. Fig. 1). The new sequence is identical to the previous one with few exceptions. In particular, a mutation in the sequence between *datZ* and *purZ* genes prevented the identification of a gene whose product corresponds to a MazG-like phosphohydrolase, named hereafter *mazZ*. We found no Shine-Dalgarno (SD) motifs upstream the genes, which is consistent with low SD motif conservation in cyanobacteria and their phages (12). Interestingly, the S-2L genome can be divided into two parts, where almost all genes follow only one direction of translation; in-between, we identified a *marR-*type repressor found typically on such junctions (13). We found 11 complete phage sequences having both *datZ* and *purZ* gene homologues (Fig. 1B), all from *Spihoviridae* family. We inferred their phylogeny and distributed all phages into four major clusters with similar genome organization; interestingly, among them cyanophage S-2L has a unique replication module. Finally, we observed that the *mazZ* gene is always co-conserved between *datZ* and *purZ*, although in one of two possible variants: MazZ-1 (S-2L-like) and MazZ-2 (ϕVC8-like). MazZ-1 undergoes frequent N-terminal fusions with other domains, but not in S-2L. In the following, we define the closely co-varying set of genes *datZ*, *mazZ* and *purZ* as the Z-cluster.

### S-2L PurZ can function as a dATPase, with no structural selection on 2’-OH

Structural analysis of ϕVC8 PurZ (PDB ID: 6FM1) suggested no particular selection on the 2’-OH group of bound ATP; thus, we sought to compare the reaction time-course of PurZ with either dATP or ATP as the phosphate donor (Suppl. Fig. S2). The HPLC result with the purified S-2L protein shows no specificity of the enzyme towards the *ribo-* and *deoxy-* variants: it can act as a dATPase but stays functional after dATP pool depletion.

We crystallized S-2L PurZ with two of its substrates, dGMP and dATP, and solved its structure at 1.7 Å resolution (Suppl. Table S1, Fig. 2B). Both ligands have identical binding mode as previously described for ϕVC8 PurZ with dGMP and ATP (PDB ID: 6FM1) (Fig. 2C).

**Figure 2.**
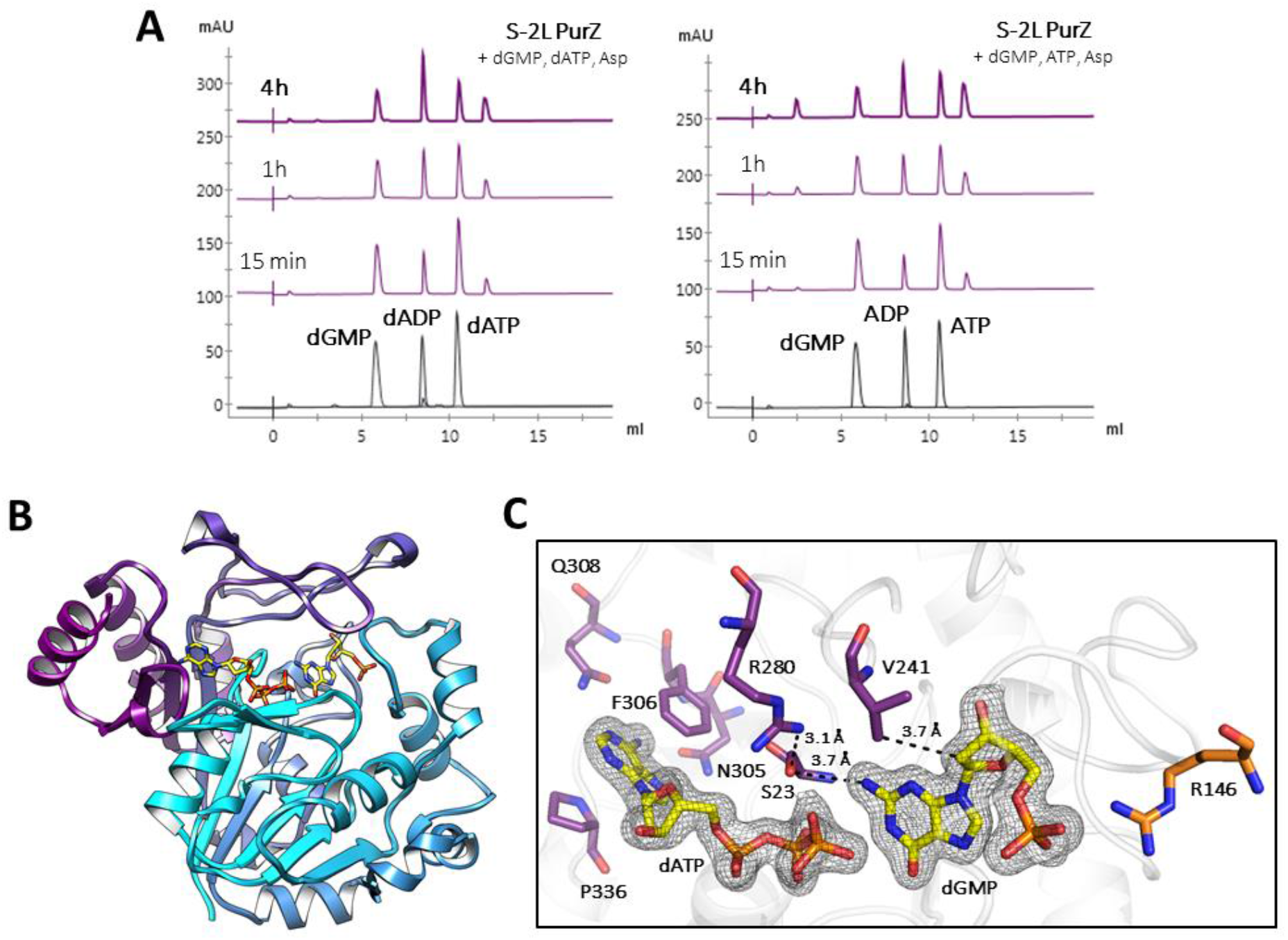
dATPase activity of S-2L PurZ: functional assay and ligand-bound structure. **A.** Time-course of the reaction catalyzed by PurZ in the presence of dGMP, Asp and dATP/ATP (purple), along with pure compounds (grey). The enzyme generates products corresponding to (d)ADP and dSMP. **B.** Ribbon representation of a PurZ crystallographic monomer, with dGMP and dATP shown in stick (yellow). **C.** Catalytic pocket of PurZ with the 2F_o_-F_c_ electron density contoured around the reactants at 1 sigma (black mesh). Crucial residues surrounding the nucleotides are coloured by chain (purple and orange).

PurA enzymes are known to be functional dimers (14, 15) or even tetramers (16). Through crystallographic symmetry, a PurZ dimer can be reconstructed for both S-2L and ϕVC8 (Suppl. Fig. 3A). Very low B-factors (Suppl. Fig. 3B) suggest an overall high rigidity for the dimer; the only exception is the Y272-L278 loop above the reactants, in contact with the aspartate substrate (Sleiman *et al*., in press). Its crucial tip (T273-T276) is strictly conserved between S-2L and ϕVC8, both in sequence and structure.

#### dGMP-binding site

Residues G22, S23, N49, A50, T132, Q190, V241 and the backbone atoms of Y21 form a pocket where the base moiety of dGMP is placed. Y21-S23 are positioned next to the amino group of dGMP in position 2. The sidechain of S23 is involved in the specificity towards the guanine base (Sleiman *et al*., in press) and its partial negative charge is stabilized by the close R280 sidechain. Residues G130, S131, R146, Y203, C204, T205 and R240 complete the dGMP pocket. Additionally, V241 is ideally placed to sterically interfere with the ribose 2’-OH group of ribonucleotides (see Discussion). Strikingly, R146 from the dimer’s other molecule forms an ion pair with the α-phosphate (17).

#### (d)ATP -binding site

Like ATP in ϕVC8 PurZ, the base moiety of dATP is stabilized by stacking interactions with F306 and P336 residues. Oxygen atoms from the side-chains of N305, Q308 and the backbone of G335 form hydrogen bonds with adenine’s 6-amino group. Importantly, the amino group of Q308 (N297 in ϕVC8) would interfere with the 2-amino group of bases G and Z. The rest of the ligand interacts with S-2L PurZ almost exclusively through its triphosphate tail with residues S23, G25, G51, H52 and T53; T53 also touches the C3’ atom side. The position inferred for the 2’-OH of ATP would be entirely exposed to the solvent, like in ϕVC8 PurZ.

### A (d)GTP-specific S-2L MazZ, an NTP-PPase with a MazG-HisE fold

Purified MazZ of cyanophage S-2L is capable of removing two terminal phosphates from a nucleotide triphosphate (Fig. 3A). Its preferential substrates are dGTP and GTP, rapidly dephosphorylated to dGMP and GMP, respectively. In contrast, S-2L MazZ shows no substantial activity with other deoxynucleotides, including dZTP (Suppl. Fig. S4).

We obtained crystals of MazZ and solved its structure, bound to the dephosphorylation product of dGTP and catalytic Mn^2+^ ions, at 1.43 Å resolution (Suppl. Table S1, Fig. 3B). Contrary to other members of the all-α NTP-PPase superfamily (18), each MazZ protein chain has two additional β-strands on its C terminus. Together, they form a homotetramer with a typical MazG-HisE fold (Suppl. Fig. S5): to our knowledge, it is the first time that the fold is recognized as identical for all MazG, MazG-like and HisE enzymes, despite the number of determined structures. The whole MazZ tetramer constitutes the asymmetric unit. The only noticeable differences in electron density between superposed chains of the tetramer lie in the solvent-exposed D43-H46 flexible loop and on both N- and C-termini.

**Figure 3.**
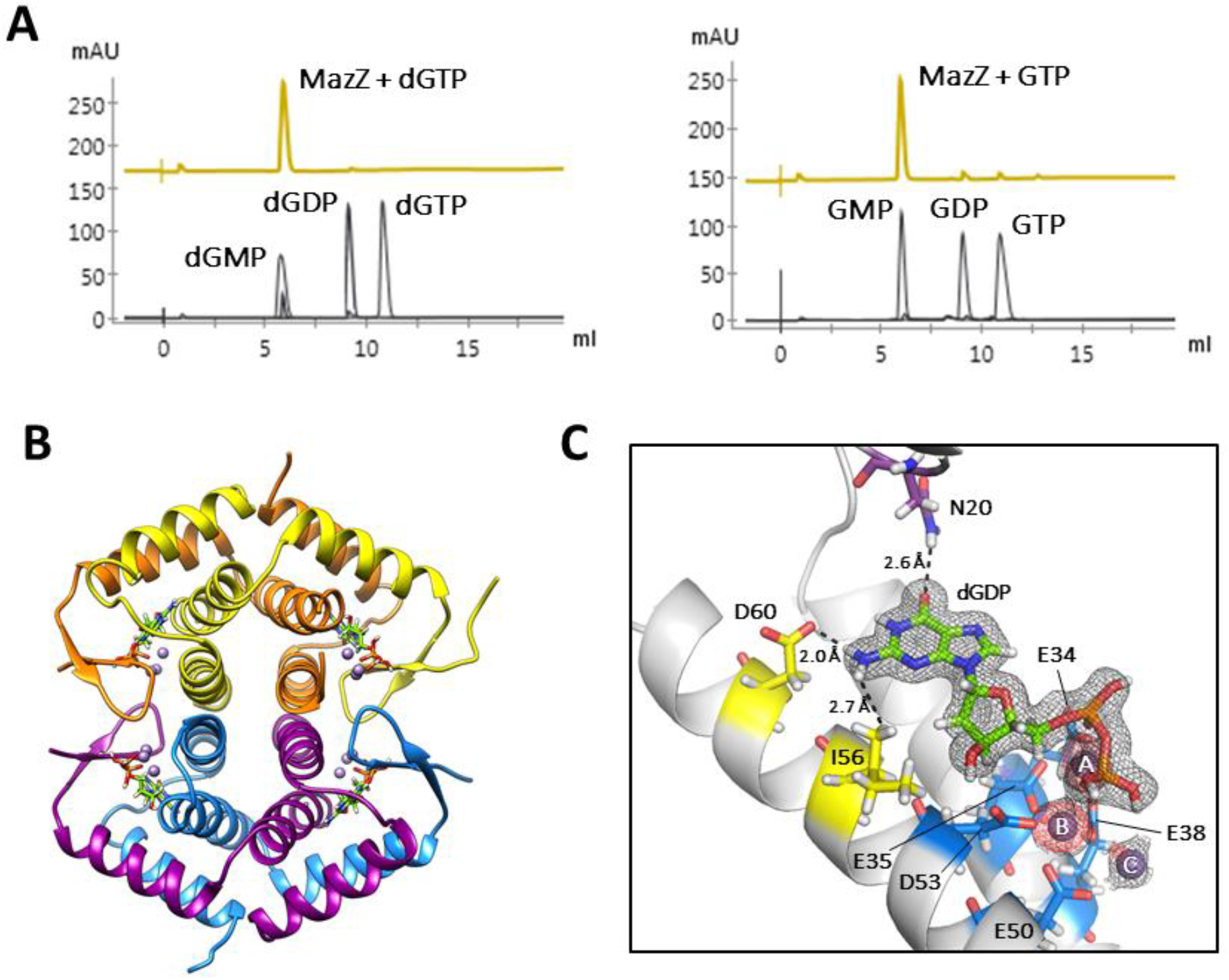
Function and structure of S-2L MazZ, a (d)GTP phosphohydrolase. **A.** HPLC analysis of S-2L MazZ activity with dGTP and GTP (gold), with nucleotide standards (black). **B.** A tetramer of MazZ, that constitutes the crystallographic asymmetric unit. Each catalytic pocket contains the reactant (lime) and three catalytic ions and results from interactions at the interface of two chains of a tight dimer. **C.** Close-up on the catalytic pocket. The product of dGTP dephosphorylation (lime) is identified as dGDP in the crystal, next to three catalytic Mn^2+^ ions (lilac spheres). The determinants of guanine specificity (yellow for N2 and purple for O6) and residues coordinating the ions (blue) are placed on a single protein chain. The 2Fo-Fc electron density around the ligands is contoured at 1 sigma (black mesh); the anomalous signal attesting for the presence of Mn^2+^ ions is contoured at 3 sigmas (red mesh).

The electron density of the dGTP dephosphorylation product revealed the presence of two phosphate groups (Fig. 3C). The dGDP intermediate observed *in crystallo* is almost completely buried inside the enzyme, except from the solvent-exposed β-phosphate side, where we found three catalytic Mn^2+^ ions.

#### dGDP -binding site

From one side, the ligand is held by residues I12, W15, I16, N20, K31, E35, D53, I56, L57 and D60 of one chain. The second chain completes the pocket with residues K76, N80, W85, A92, M93, R94 and H95. The closest residue to guanine’s O6 is N20, through its amide nitrogen at 3.2 Å; the 2-amino group is only 3.0 Å away from the carboxyl group of D60. Altogether, the specificity of MazZ emerges from the cavity volume matching guanine’s shape and its charge compatibility.

We observe no steric hindrance for the possible 2’-OH group. Mutation of the closest I56 could affect the specificity for ribonucleotides; however, it also contacts the base’s 2-amino group and is surrounded by an intricate network of other residues.

#### Catalytic Mn^2+^ ions

The three Mn^2+^ ions, named A, B and C, are strictly equivalent to the Mg^2+^ ions found in a related NTP-PPase (19). The ion A is coordinated by residues E34, E35 and E38; ion B by the E50 and D53; and ion C by E38 and E50. These ions are all coordinated in an octahedral fashion with protein, ligand and water atoms. It is possible that the two-metal-ion mechanism described for NTP-PPases occurs in fact with two consecutive dephosphorylation steps, justifying the presence of the three ions and intermediate dGDP. Additionally, residue R83, positioned only 2.8 Å away from the β-phosphate, is probably important for stabilizing the transition state (20).

### Successful conversion of *E. coli* DNA with S-2L’s Z-cluster

We now propose a complete 2-aminoadenine metabolic pathway for cyanophage S-2L and related *Siphoviridae* phages (Fig. 4A). The post-infection composition of cellular dNTP pool is highly modified: dATP is eliminated and replaced by dZTP, which is created in turn from dGTP. In the case of cyanophage S-2L, the non-specific Primase-Polymerase (not conserved in other *Siphoviridae* possessing the Z-cluster) readily inserts 2-aminoadenine in front of any instructing thymine.

**Figure 4.**
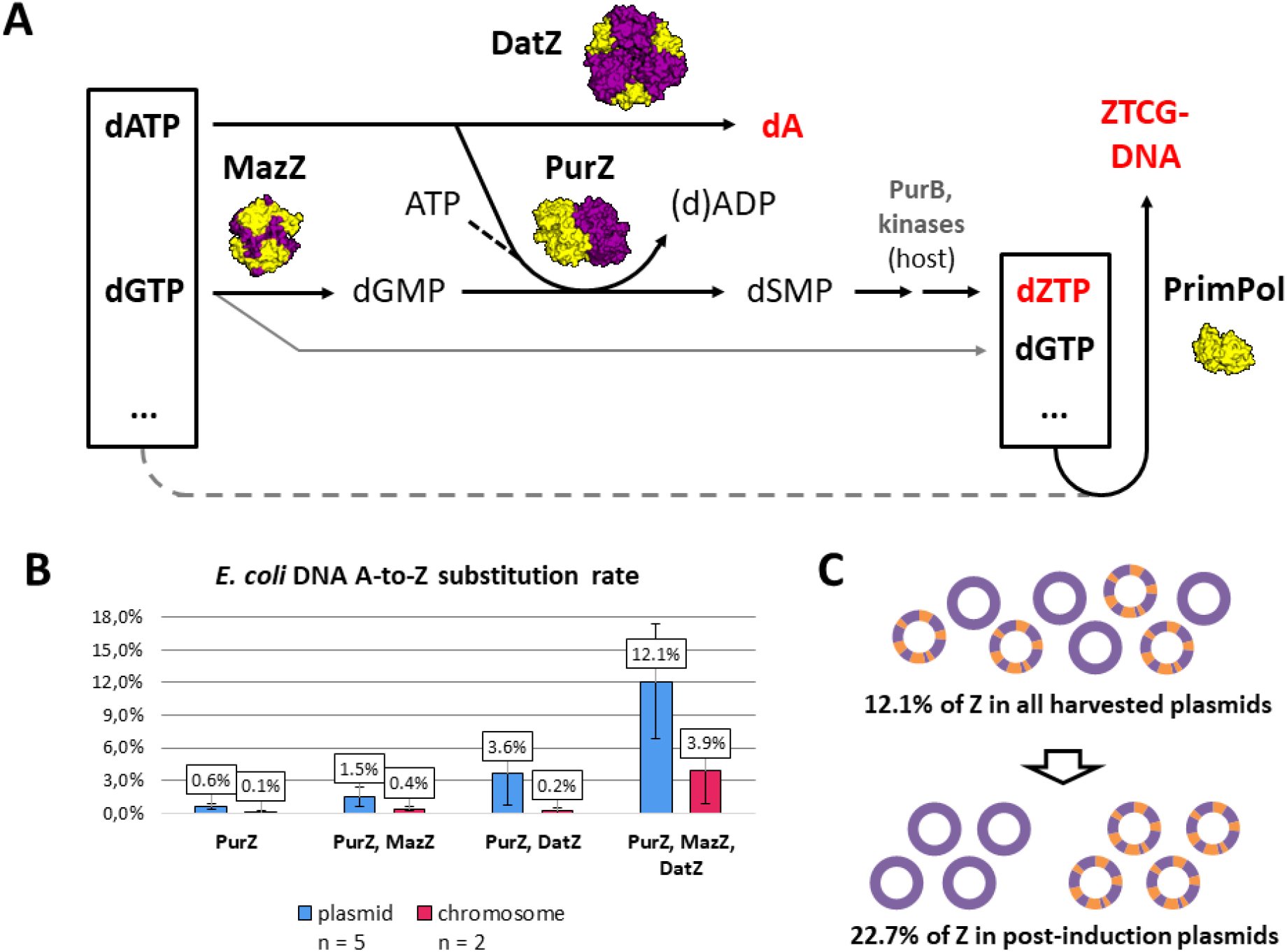
Metabolic pathway of 2-aminoadenine in S-2L phage and result of its introduction in *E. coli*. **A.** The cellular original pool of nucleotides is represented by the box to the left, the one modified through expression of the viral Z-cluster is shown on the right. Three dots represent the unmodified dTTP and dCTP. The structures of the involved S-2L proteins solved in this and a previous work (11) are shown next to their respective reaction arrows (to scale). Host enzymes (names in grey) finalise the dZTP pathway (Sleiman *et al*., in press). The thin, grey arrow stands for no modification. The dashed grey arrow stands for potential use of the standard dNTP pool by PrimPol in the absence of the Z-cluster. **B.** Average A-to-Z substitution rates in total plasmidic and chromosomal DNA obtained after complete or partial transplantation of S-2L’s Z-cluster to *E. coli* (labels below). **C.** Accounting for the pre-induction native fraction, roughly every fourth plasmidic adenine (violet) was changed to 2-aminoadenine (orange) after the induction of the Z-cluster.

We put the genes *datZ*, *mazZ* and *purZ* under the control of the T7 promoter on compatible expression plasmids in *E. coli* and induced the cultures in exponential phase. The whole system proved to be toxic to the cells when expressed (Suppl. Fig. S6). Whether or not it is true for the natural hosts of S-2L and ϕVC8 is uncertain but we note that these phages are known to be lytic anyway (7, 21).

Using mass spectrometry, we were able to quantify the Z content in bacterial DNA (Fig. 4B). In plasmids and the chromosome, cells substituted on average 12.1% and 3.9% of total adenine content with 2-aminoadenine, respectively. Additionally, by quantifying the non-modified pre-induction plasmidic fraction, we normalized the plasmidic A-to-Z substitution ratio to 22.7% after expression of the Z-cluster (Fig. 4C). Importantly, incomplete Z-cluster expression yielded lower substitution rates and the effect of *datZ* and *mazZ* absence is synergistic. Altogether, it appears that bacterial DNA can be enriched with diaminopurine efficiently with the introduction of the Z-cluster alone.

## Discussion

### Molecular basis for substrate selectivity in PurZ vs PurA

To compare PurZ and PurA, we prepared a structure-informed sequence alignment. 36 residues are strictly conserved between PurZ and PurA, while 26 further residues have very little variability (Suppl. Fig. S7). These two sets of residues cluster in the catalytic pocket and the surrounding layer, respectively (Suppl. Fig. S8A). Finally, 16 residues are unique to PurZ. Their placement is intermediate compared to the previous classes, although several such residues (S23, T273 and Q308) interact with the substrates. In PurA a conserved arginine (R303 in *E. coli*) balances the partial charge of O2’ of IMP at 2.5-4 Å and interacts with the free aspartate substrate (22). The corresponding residues in PurZ extend noticeably further, being 7.9 Å (S-2L R244) and 8.5 Å (ϕVC8 K267) away from C2’, precluding any possible stabilization of the ribonucleotide. Moreover, in PurZ an insertion deforms the loop that contains a conserved V241, pulling it slightly closer to the C2’ ribose atom – from 4.3-5.3 Å as seen in PurA to 3.7 Å (S-2L) and 3.6 Å (ϕVC8). Lastly, one of the two deletions specific to PurZ and PurA from *P. horikoshii* (Sleiman *et al*., in press) are compensated by an additional C-terminal helix α_11_ unique to S-2L (Suppl. Fig. S8B). Using the above alignment, we constructed a phylogenetic tree (Suppl. Fig. S9). Strikingly, this tree points to a possible archaeal origin for PurZ.

### S-2L MazZ as a representative of MazZ-1 and similarity with MazZ-2

With the exception of E34, all residues of S-2L MazZ coordinating the catalytic ions are strictly conserved across all MazZ-1 homologues (Suppl. Fig. S10). Most of the amino acids participating in nucleotide pocket formation tend to be conserved as well: only the residues I16, N20 and A92 placed around the nucleobase are unique to S-2L. Importantly, residues I56 and D60 are kept by all MazZ-1 homologues, suggesting preserved guanine specificity. In MazZ-2, 4 negatively charged residues are also strictly conserved, their position corresponding to E35, E38, E50 and E53 of S-2L MazZ (Suppl. Fig. S11). However, other nucleobase-binding residues differ from MazZ-1 consensus, implying alternative binding mode of the substrate. Finally, a counterpart of R83 important for the two-metal-ion mechanism is preserved in both variants of MazZ.

### 2-aminoadenine metabolic pathway and ZTGC-DNA

On the day of submitting this paper, we became aware of the work of Zhou et al. (23) on the same subject. These authors identify 3 genes conserved in several phages that contain Z in their genome: *purZ*, also described by one of us and his collaborators (24), a dATPase, which is similar to the datZ gene that we described recently (11) and a gene called DUF550, which is both a dATPase and a dGTPase, found in the ϕVC8 phage, that provides the dGMP substrate for the reaction catalyzed by PurZ. This gene is similar to (but longer than) the *mazZ* gene that we describe here in phage S-2L, and corresponds to the mazZ-2 variant as shown in Figure 1B. In ϕVC8 it is apparently both a dATPase and a dGTPase but in S-2L it is only a dGTPase, which underlines some flexibility in the strategy of these phages to incorporate Z instead of A in their genome. Another variation in this strategy comes from the DNA polymerases of these phages: all of them but S-2L have a polA (pol I) DNA polymerase, that was shown recently by Pezo et al. (25) to have some specificity of incorporation of Z vs A in front of T, whereas S-2L has only a PrimPol DNA polymerase, which has no such specificity (11).

Pre- and post-replicative modification pathways of genomic thymidine in bacteriophages have already been successfully transplanted to *E. coli* (26) or found to be active in its lysates (27). However, these changes involved groups protruding on the outside of the double helix major groove, as in epigenetic modifications. In the case of 2-aminoadenine, the additional amino group on the Watson-Crick edge profoundly modifies the stability of base-pairs between the two nucleic acid strands, changing the thermodynamics of the DNA molecule rather than its external accessibility.

The kinetics of creating dZTP from dGTP has to be fine-tuned by the infecting viruses, as dGTP is also a crucial substrate for the synthesis of ZTGC-DNA. We note that in *in vitro* assays S-2L PurZ is overall less active than the homologous *E. coli* PurA (Fig. 2A), which may contribute to this effect.

In addition, we show here that the Klenow fragment of *E. coli* Pol I, a DNA polymerase participating in plasmid replication (28), does not discriminate between A and Z whatsoever (Suppl. Fig. S12). This relaxed specificity holds true for most DNA and RNA polymerases (29).

The toxic effect of 2-aminoadenine in ATGC-DNA organisms such as *E. coli* probably stems from the complexity of the regulation of DNA transactions: sigma factors are sensitive to A-to-Z substitutions (30), while replication origins are known to require AT-rich sequences where melting is facilitated (31). Nonetheless, the melting point of a Z:T homopolymer was found to be intermediate between the A:T and G:C ones (32). Attuning the cellular machinery for the presence of 2-aminoadenine should thus be in principle achievable, as it is possible for the less complex phages, leading potentially to a pure ZTGC-DNA synthetic organism.

## Acknowledgements

We thank the Crystallogenesis and Crystallography Platform (PF6) of Institut Pasteur for help in crystallization and preliminary crystallographic data collection. We thank M. Hollenstein and his group for the use of their HPLC system. We also thank staff from PROXIMA-1 and PROXIMA-2 beamlines for help in data collection and SOLEIL (Saint-Aubin, France) and for provision of synchrotron radiation facilities. We thank P. Legrand for discussions regarding the structural aspect of this work.

## Author contributions

MD directed the study. DC and MD designed the study. DC, FB and PAK performed experiments. DC, FB, PAK and MD analysed the data. DC and MD wrote the manuscript. All authors edited the manuscript.

## Data availability

The data that support the findings of this study are available from the corresponding author upon request.

The x-ray structures described in this paper have been deposited to the PDB under the codes of 7ODX and 7ODY along with the diffraction data.

## Materials and methods

### S-2L genome sequencing, annotation and homology with other *Siphoviridae*

S-2L DNA was isolated from phage lysate of *Synechococcus elongatus* culture, adapting the techniques commonly used for phage λ DNA. The S-2L genomic library was prepared by successive DNA fragmentation, adapter ligation and amplification by GATC Biotech. Libraries were sequenced using Illumina HiSeq. 14,198,980 sequenced reads (2 x 150 bp) were obtained, covering 4,259,694,000 bases. Resulting reads were mapped against the GenBank AX955019.1 S-2L reference sequence using Minimap2 v2.17 (33). Variant calling was carried out by Freebayes v1.3.2 (34) and later filtered by VCFLIB (35). A consensus sequence was generated using VCF Consensus Builder v0.1.0 (36). Annotation of the consensus sequence was carried out by translated BLAST (37). Representation of the S-2L genome was made with SnapGene Viewer (38). Phages related to S-2L through *purZ* and *datZ* genes were found by homology searches using NCBI BLAST (37), with protein sequences as separate queries.

Variants MazZ-1 and MazZ-2 correspond to two sets of MazG-like family proteins whose genes are co-conserved with *purZ* and *datZ.* In each set, they have an average 44% (±6) and 43% (±9) protein sequence identity; BLAST does not find any significant hits between the two sets. Distant identity of 29-31% is observed only for Ochrophage vB_OspB_OH MazZ-2 and additional N-terminal domains of Salmophage PMBT28 and Proviphage Kokobel2 fused to C-terminal MazZ-1 sequences. Difference between MazZ-1 and MazZ-2 is further highlighted by divergent conservation profiles presented in Supplementary Data.

### Protein expression and purification

Synthetic genes for expressed proteins were optimized for *E. coli* and synthesized using ThermoFisher’s GeneArt service. Genes were cloned into modified pRSF1-Duet expression vector with a TEV-cleavable N-terminal 14-histidine tag (39) using New England Biolabs and Anza (Thermo Fisher Scientific) enzymes. *E. coli* BL21-CodonPlus (DE3)-RIPL cells (Agilent) were separately transformed with engineered plasmids. Bacteria were cultivated at 37°C in LB medium with appropriate antibiotic selection (kanamycin and chloramphenicol), and induced at OD = 0.6-1.0 with 0.5 mM IPTG. After incubation overnight at 20°C, cells were harvested and homogenized in suspension buffer: 50 mM Tris-HCl pH 8, 400 mM NaCl, 5 mM imidazole. After sonication and centrifugation of bacterial debris, corresponding lysate supernatants were supplemented with Benzonase (Sigma-Aldrich) and protease inhibitor (Thermo Fisher Scientific), 1 μl and 1 tablet per 50 ml, respectively. Proteins of interest were isolated by purification of the lysate on Ni-NTA column (suspension buffer as washing buffer, 500 mM imidazole in elution buffer). Histidine tags were removed from the proteins by incubation with His-tagged TEV enzyme overnight. After removing TEV on Ni-NTA column, proteins were further purified on Superdex 200 10/300 column with 25 mM Tris-HCl pH 8, 300 mM NaCl. All purification columns were from Life Sciences. Protein purity was assessed on an SDS gel (BioRad). The enzymes were concentrated to 10-19.5 mg ml^−1^ with Amicon Ultra 10k and 30k MWCO centrifugal filters (Merck), flash frozen in liquid nitrogen and stored directly at −80°C, with no glycerol added.

### Nucleotide HPLC analysis

30 µM (1.25 mg ml^−1^) of S-2L PurZ or 4.2 µM (0.2 mg ml^−1^) of *E. coli* PurA was incubated at 37°C for 1h (if not stated otherwise) with 2 mM of respective nucleotides and 10 mM of aspartate, in a buffer containing 50 mM Tris pH 7.5 and 5 mM MgSO_4_. For S-2L MazZ, 10 μM of the enzyme was incubated at 37°C for 15 min with 100 μM of respective nucleotides, in a buffer containing 50 mM Tris pH 7.5 and 5 mM MgCl_2_. Reaction products were separated from the protein using 10 000 MWCO Vivaspin-500 centrifugal concentrators and stored in −20°C. Products and standards were assayed separately, using around 40 nmol of each for anion-exchange HPLC on DNA-PAC100 (4×50 mm) column (Thermo Fisher Scientific). After equilibration with 150 μl of a suspension buffer (25 mM Tris-HCl pH 8, 0.5% acetonitrile), nucleotides were injected on the column and eluted with 3 min of isocratic flow of the suspension buffer followed by a linear gradient of 0-200 mM NH_4_Cl over 12 min (1ml min^−1^). Eluted nucleotides were detected by absorbance at 260 nm, measured in arbitrary units [mAu]. High-purity nucleotides and chemicals were bought from Sigma Aldrich, and HPLC-quality acetonitrile was from Serva.

### Transplantation of the Z-cluster into *E. coli* and 2-aminoadenine detection

Gene *datZ* was cloned onto plasmid pET100/D-TOPO providing ampicillin resistance. Genes *purZ* and *mazZ* were cloned into modified pRSF1-Duet providing kanamycin resistance, in slots 1 (his-tagged) and 2, respectively, each with a ribosome-binding site upstream. Constructs with partial Z-cluster did not include *mazZ* in pRSF1-Duet slot 2 (*datZ*/*purZ* or *purZ* alone) and further had *datZ* replaced by *mazZ* on pET100/D-TOPO (*mazZ*/*purZ*). One or both plasmids were used to transform *E. coli* BL21-CodonPlus (DE3)-RIPL cells.

Bacteria were cultivated at 37°C in LB medium with appropriate antibiotic selection and induced with 1 mM IPTG at OD = 0.60 (± 0.03). After 2 hours, plasmids were isolated with NucleoSpin Plasmid QuickPure kit (Macherey-Nagel) and suspended in water. When not induced, natural leaking of the promotors resulted in 0.01% of Z content, which was not lethal to the cells. Additionally, we observed that the expression of DatZ lowers the overall plasmidic DNA yield, that can be partially compensated when both MazZ and PurZ are expressed as well; this is in line with the proposed 2-aminoadenine metabolism pathway.

Chromosomal and plasmidic DNA was digested to nucleosides with Nucleoside Digestion Mix (NEB) and separated on Amicon Ultra-0.5 mL 10K centrifugal filters. Nucleosides were analysed by LCMS, with standard nucleoside controls (Sigma Aldrich) or dZ (Biosynth Carbosynth). dZ was found to elute at the same position than dG, but with strikingly different MS/MS profiles.

The A-to-Z substitution rate was taken as a ratio of dZ content to dZ+dA. For plasmids, this number was further normalized to the post-induction fraction by calculating the ratio between the pre-induction and final plasmidic DNA yield.

### Crystallography and structural analysis

All crystallization conditions were screened using the sitting drop technique on an automated crystallography platform (40) and were reproduced manually using the hanging drop method with ratios of protein to well solution ranging from 1:2 to 2:1. PurZ was screened at 19 mg ml^−1^ with a molar excess of 1.2 of dGMP in 4°C. Capped thick rods grew over several days in 15% v/v Tacsimate and 2% w/v PEG 3350 buffered with 100 mM HEPES pH 7, and did not appear in the absence of dGMP. MazZ was screened at 14.7 mg ml^−1^ with a molar excess of 1.2 of dGTP at 18°C. Big bundles of thin needles grew over a week in 200 mM Li_2_SO_4_ and 1.26 M (NH_4_)_2_SO_4_ buffered with 100 mM Tris pH 8.5. PurZ crystals were soaked for 15 min in a solution containing 30% glycerol, 50% dATP solution (100 mM) and 20% crystallization buffer; MazZ crystals were soaked for several seconds with 30% glycerol and 70% crystallization buffer, with added 30 mM MnCl_2_. All crystals were then frozen in liquid nitrogen. Crystallographic data was collected at the Soleil synchrotron in France (beamlines PX1 and PX2), processed by XDS (41) with the XDSME (42) pipeline and refined in Phenix (43). Nucleotide constraints for structure refinement were obtained using Grade Web Server (44). The structure of S-2L PurZ was solved by molecular replacement with ϕVC8 PurZ model (PDB ID: 6FM1). The structure of MazZ was solved using anomalous signal from bound Mn^2+^ ions that guided automatic model-building in Phenix’ AutoSol, with final manual reconstruction steps using Coot (45). PurZ quaternary structure analysis was performed using PISA (46) and suggested the presence of a stable dimer. The presence of two phosphate groups in dGTP dephosphorylation product bound to MazZ was confirmed by a careful analysis of electronic density and B-factors, eliminating the possibility of a flexible γ-phosphate. Density corresponding to dGDP was also observed in a structure from a crystal not soaked in MnCl_2_. HPLC analysis of dGTP substrate in solution excluded dGDP contamination.

### DNA polymerase assay

Fluorescence-based polymerase activity test was executed in 20 mM Tris-HCl pH 7 and 5 mM MgCl_2_, with 3 μM of dT_24_ overhang DNA template, 1 μM of FAM 5’-labeled DNA primer and various concentrations of either dATP or dZTP. The Klenow polymerase was added to final concentration of 5 U in 50 μl. The assay was conducted at 37°C for 5 min. Before adding the protein, DNA was hybridized by heating up to 95°C and gradually cooling to reaction temperature. Reactions were terminated by adding two volumes of a buffer containing 10 mM EDTA, 98% formamide, 0.1% xylene cyanol and 0.1% bromophenol blue, and stored in 4°C. Products were preheated at 95°C for 10 min, before being separated with polyacrylamide gel electrophoresis and visualised by FAM fluorescence on Typhoon FLA 9000 imager. All oligonucleotides were from Eurogentec, chemicals from Sigma-Aldrich, Klenow polymerase from Takara Bio, dATP from Fermentas (Thermo Fisher Scientific) and dZTP from TriLink BioTechnologies.

### Structure and sequence alignments, phylogeny

Structures homologous to PurZ available in PDB were identified using Dali server (47). The sequences were aligned in PROMALS3D (48) using structural data supplemented by full protein sequences. Distances in PurA molecules were measured using PDB IDs 1P9B, 5I34, 5K7X (Arg-O2’ of IMP) and 1P9B, 2DGN, 2GCQ, 4M9D, 5I34, 5K7X (Val-C2’); for ϕVC8 PurZ measurements, PDB ID 6FKO was used. Graphical multialignments were prepared with ESPript 3 (49). Maximum-likelihood phylogenetic tree was prepared with MEGA X (50) with default parameters, with 100 bootstrap replications. Protein structures were analysed with Chimera (51) or Pymol (52).

