## Supplementary material for "Characterization of a triad of genes in cyanophage S-2L sufficient to replace adenine by 2-aminoadenine in bacterial DNA"

### Supplementary Figures

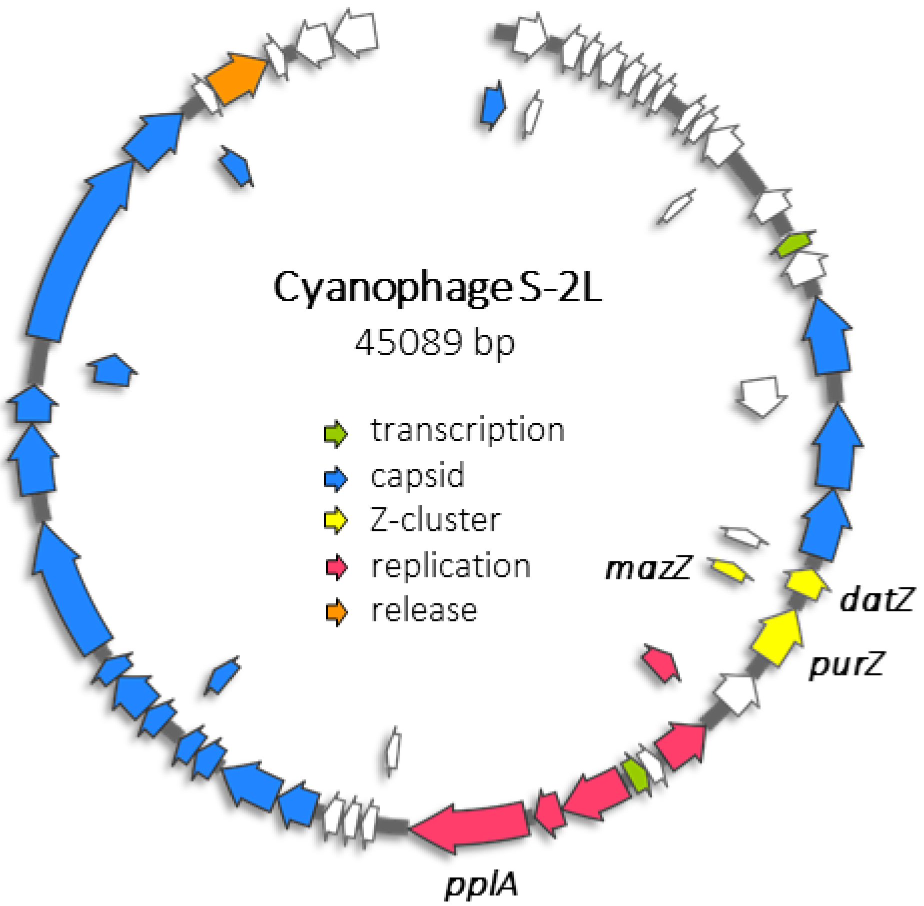


**Supplementary Figure S1.** Map of S-2L genome with all identified genes (arrows) coloured according to their function, as indicated inside the circle. The names of the genes of interest (Z-cluster), with the addition of the gene of PrimPol (*pplA*), are indicated to visualize their positions.

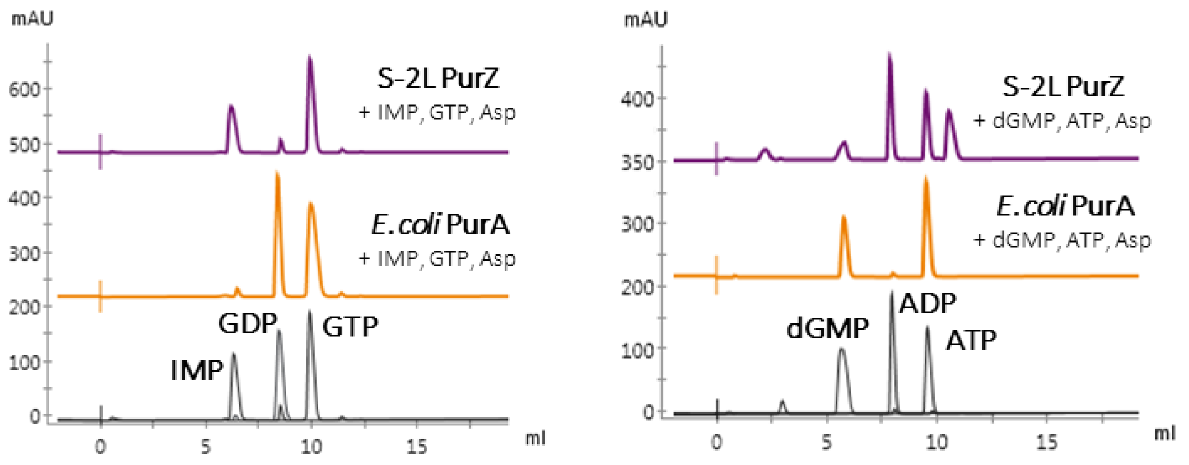


**Supplementary Figure S2. Comparison of the enzymatic activities of PurZ of S-2L with PurA of *E. coli***. In the reaction time of 15 min, PurA rapidly transforms IMP, GTP and Asp mixture into GDP and sIMP, whereas PurZ stays inactive even at higher concentration (left panel). Inversely, PurZ catalyses the reaction from dGMP, ATP and Asp to ADP and dSMP, contrary to PurA that does not recognise these substrates (right panel). Although the sIMP peak is confounded with the GTP one, it gives a noticeable shift in 260/280 nm absorbance ratio.

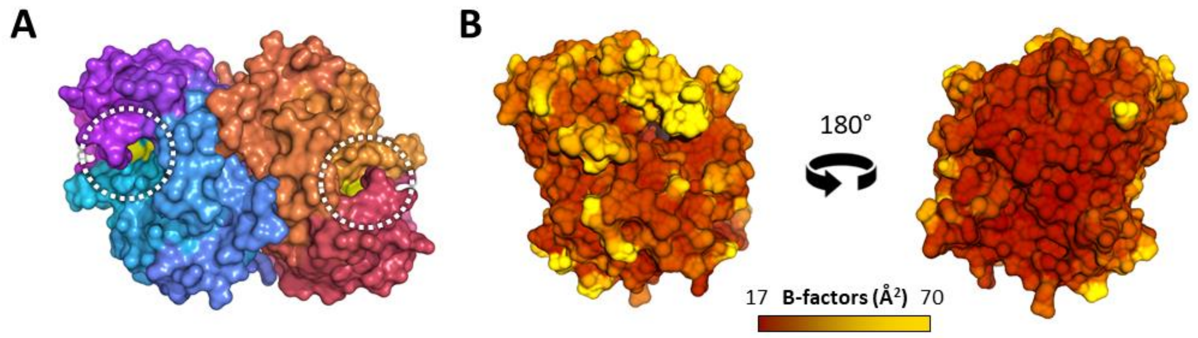


**Supplementary Figure S3. Properties of the S-2L PurZ structure highlighting its dimeric state. A.** PurZ dimer, in surface representation: the two domains are coloured in cyan-purple and orange-light red gradients. White dotted circles point to the opposite catalytic sites. **B.** Surface representation of PurZ coloured using experimental B-factors, with the corresponding scale bar below. The flexible loop above the catalytic cleft (left) define the aspartate loop. The interface between the dimer (right) is particularly rigid, strongly suggesting a constitutive dimeric form of PurZ.

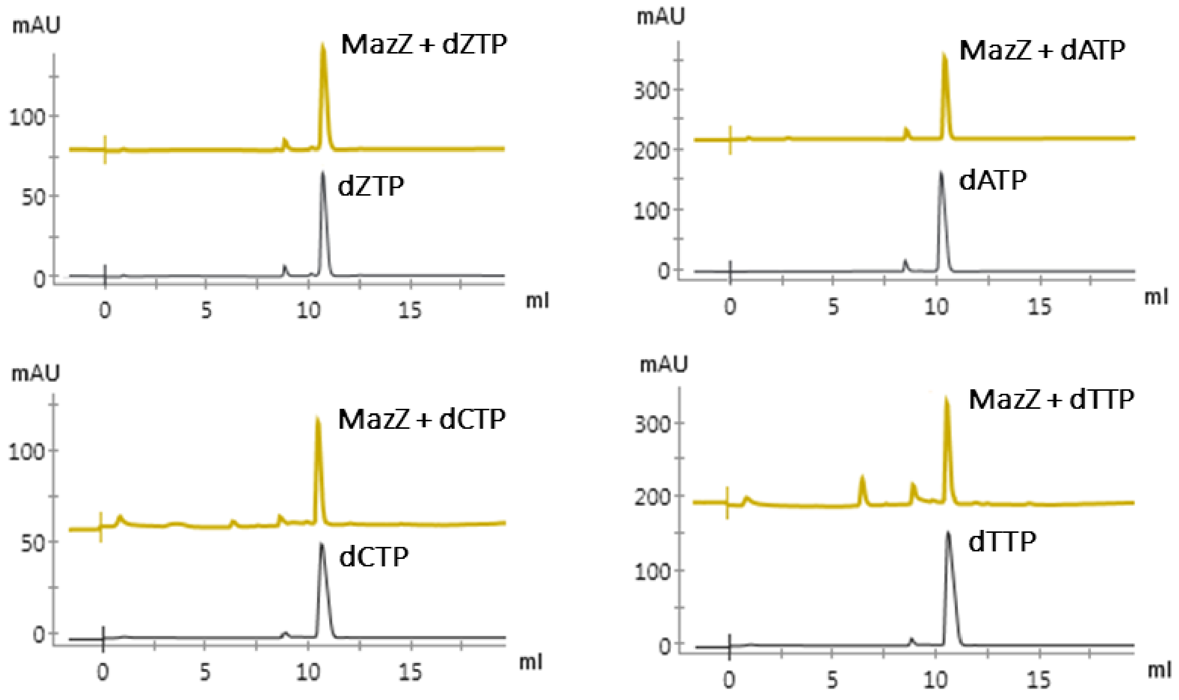


**Supplementary Figure S4. Lack of S-2L MazZ specificity for deoxynucleotide triphosphates other than dGTP, including dZTP**. Nucleotide standards are in black, the products eluted after incubation of the corresponding triphosphates with MazZ are in gold.

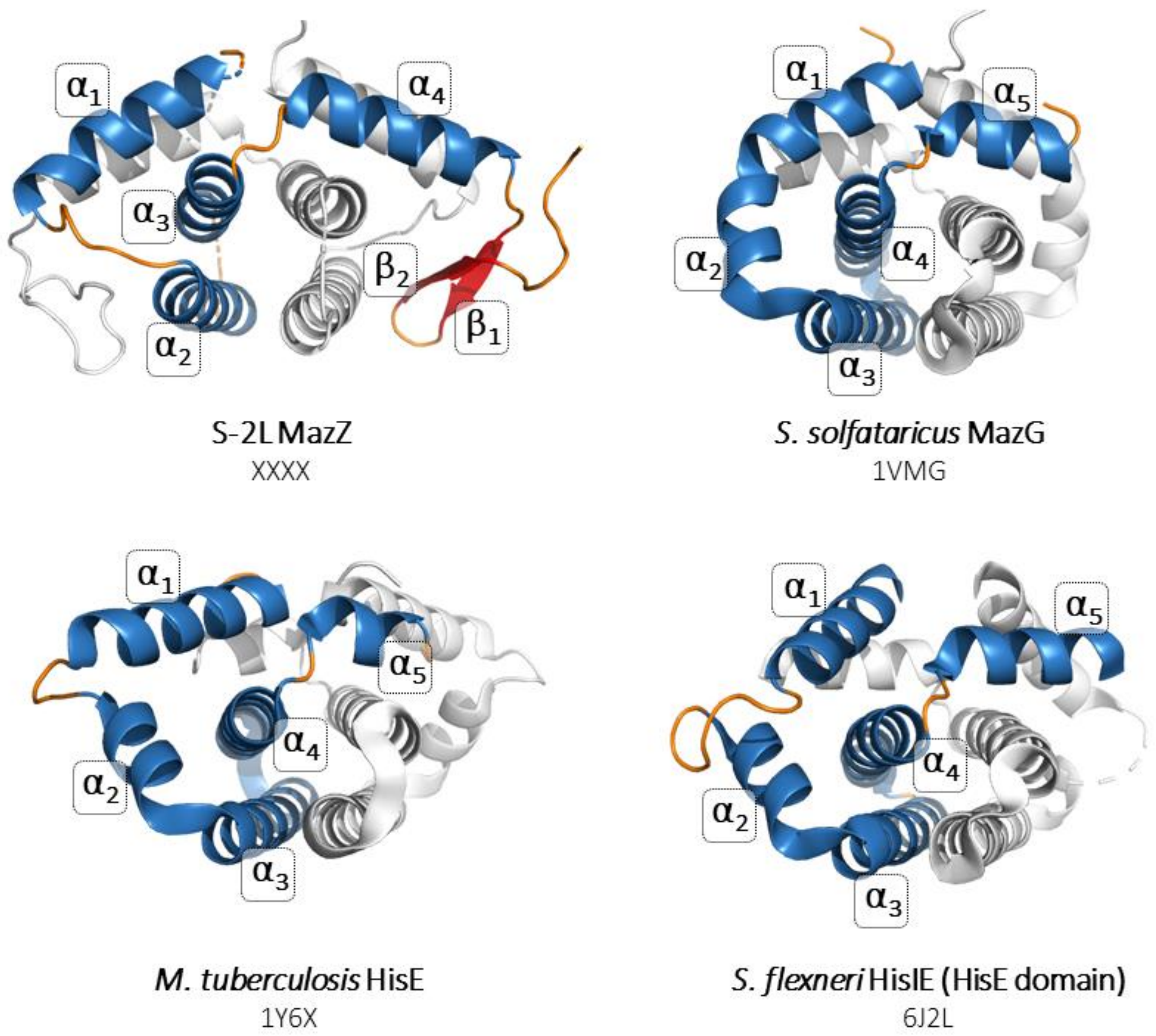


**Supplementary Figure S5. Common fold shared by S-2L MazZ, bacterial MazG and HisE proteins.** The tight dimer part is shown for four exemplary structures, viewed from the same perspective. For each enzyme both chains are shown in ribbon representation. One of the chains is coloured: α-helices in blue, loops in orange and β-strands in red. For the coloured chain, the secondary structure elements are numbered. Below each image, the organism of origin, protein name and the PDB code are indicated.

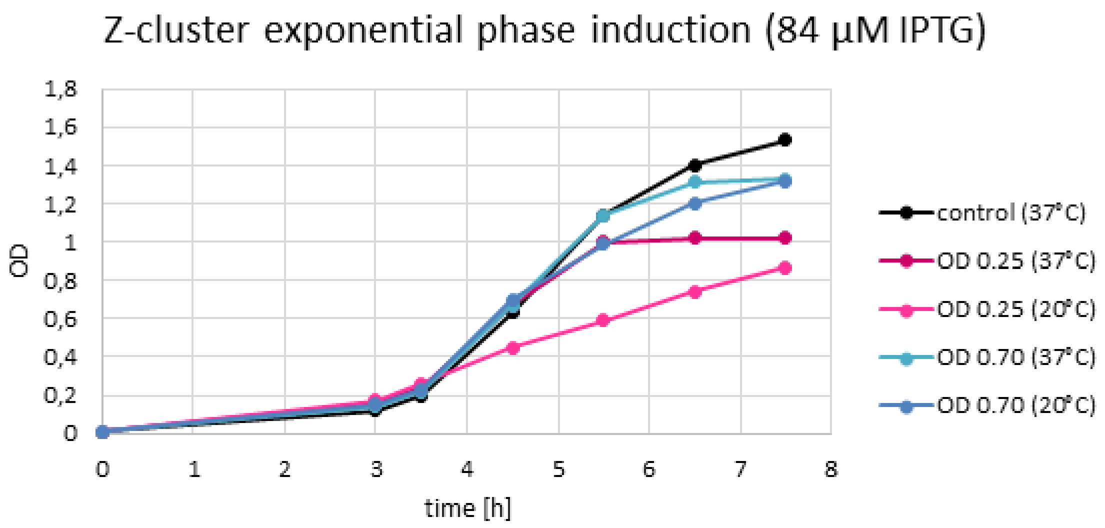


**Supplementary Figure S6. Toxic effect of the expressed Z-cluster on *E. coli*.** Cultures were induced with low concentration of IPTG at two points of the exponential growth phase and incubated in 20 or 37°C afterwards (legend to the right). All conditions showed arrested growth in a matter of a few generations compared to a non-induced control.

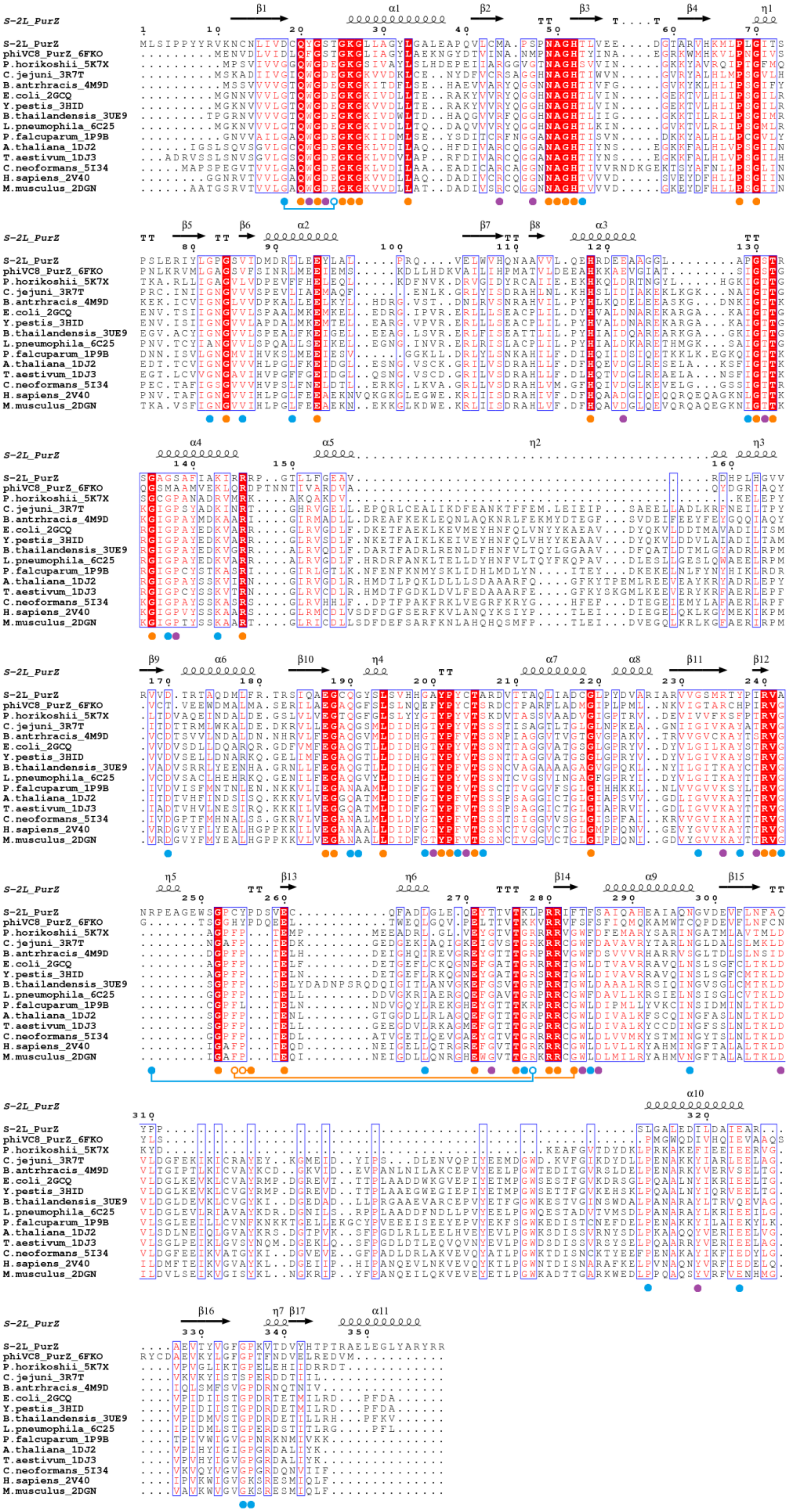
S-2L_PurZ
phiVC8_PurZ_6FKO
P.horikoshii_5K7X
C.jejuni_3R7T
B.antrhracis_4M9D
E.coli_2GCQ
Y.pestis_3HID
B.thailandensis_3UE9
L.pneumophila_6C25
P.falcuparum_1P9B
A.thaliana_1DJ2
T.aestivum_1DJ3
C.neoformans_5I34
H.sapiens_2V40
M.musculus_2DGN

**Supplementary figure S7. Structural multialignment between S-2L PurZ and all 14 homologous synthases available in the PDB**. Organism names and PDB codes are indicated on the left. Full circles below the alignment mark positions of residues of interest, divided into three categories: residues strictly conserved in all representatives (orange); loosely conserved residues with two similar variants or occasional mutations (blue); residues strictly conserved in PurA, but not in PurZ enzymes (purple). Empty circles highlight conserved residues with sequence rearrangements that have a shifted backbone position with respect to S-2L's ones (connected full circles), but with superposing functional groups. Occasional unstructured and unbuilt regions were filled with sequence information alone and no structural information.

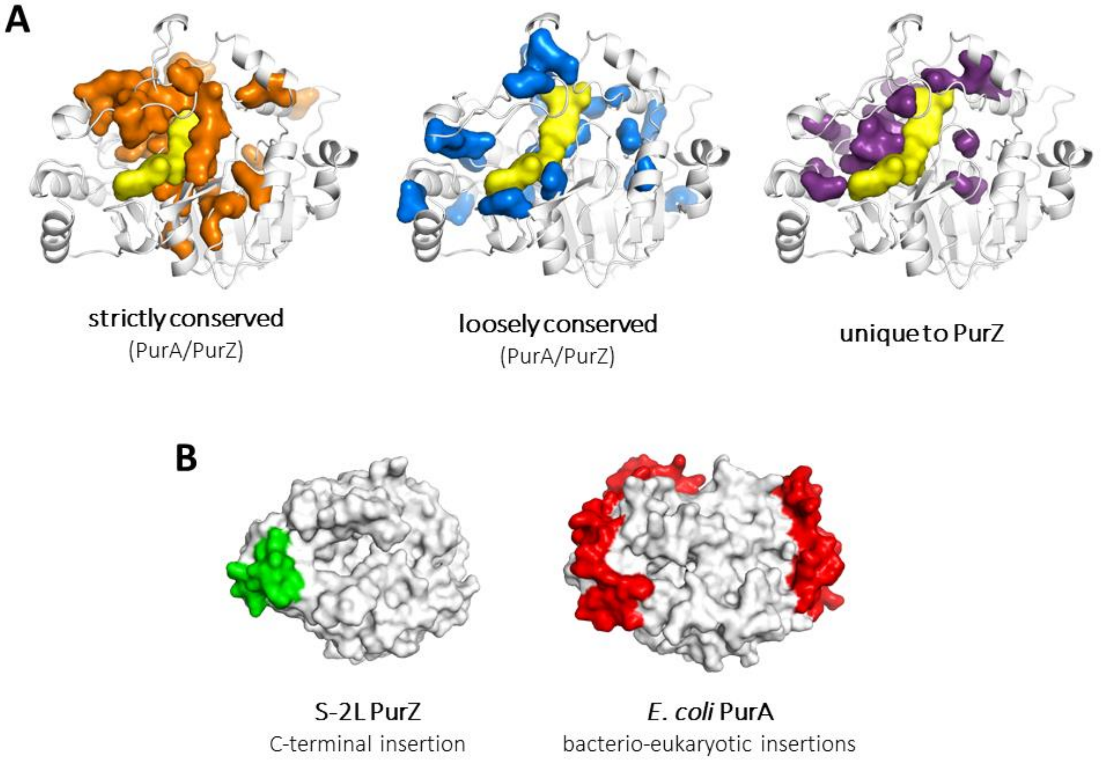


**Supplementary Figure S8. Visualisation of relationships between S-2L PurZ and its structural homologues. A.** Conserved residues from Suppl. Fig. S7 are mapped onto S-2L PurZ structure, using the same colour code. Nucleotide substrates – dGMP and dATP – are in yellow. **B.** Visualisation of a S-2L-specific C-terminal insertion in form of an alpha-helix (green, left) and bacterio-eukaryotic insertions (archaeo-viral deletions) of two large segments, shown on *E.coli* PurA (red, right).

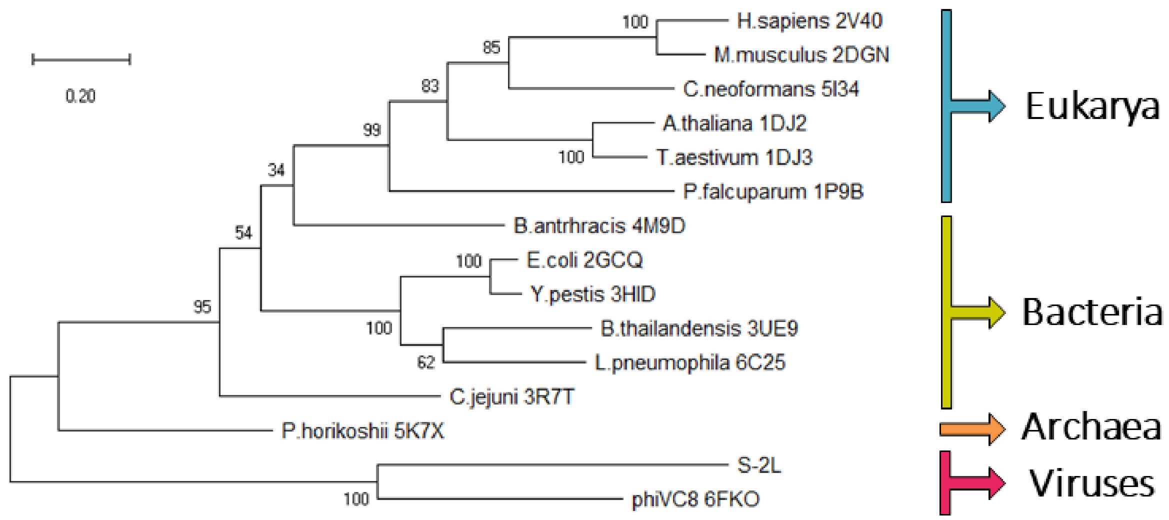


**Supplementary Figure S9. Non-rooted maximum-likelihood phylogenetic tree of PurA/PurZ synthases.** The tree was generated using the structural alignment from Fig. S7. Enzymes are divided into four clades: eukaryotic, bacterial, archaeal and viral, the two last ones sharing a recent ancestor. The reference distance corresponds to an average 0.2 substitution per site. The topology of the bootstrap consensus tree is identical, supporting the result presented here.

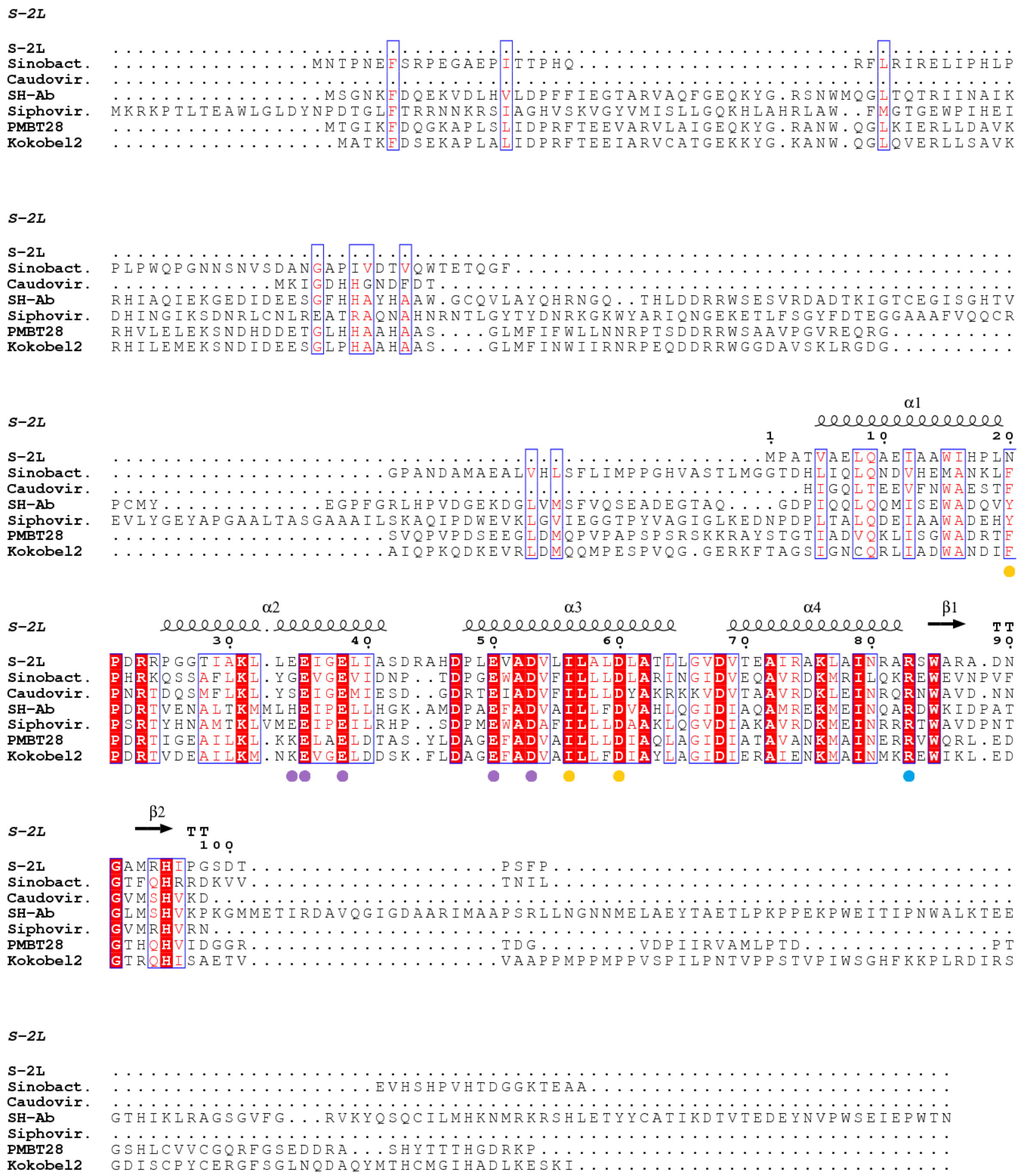


**Supplementary Figure S10. Sequence multialignment of MazZ-1 homologues (S-2L-like).** Numbering above the alignment refers to S-2L MazZ. Catalytic residues of S-2L MazZ are marked with lilac dots, R83 stabilising an intermediate product is in blue, residues coordinating 2-amino group, O6 and sugar moiety are in orange.

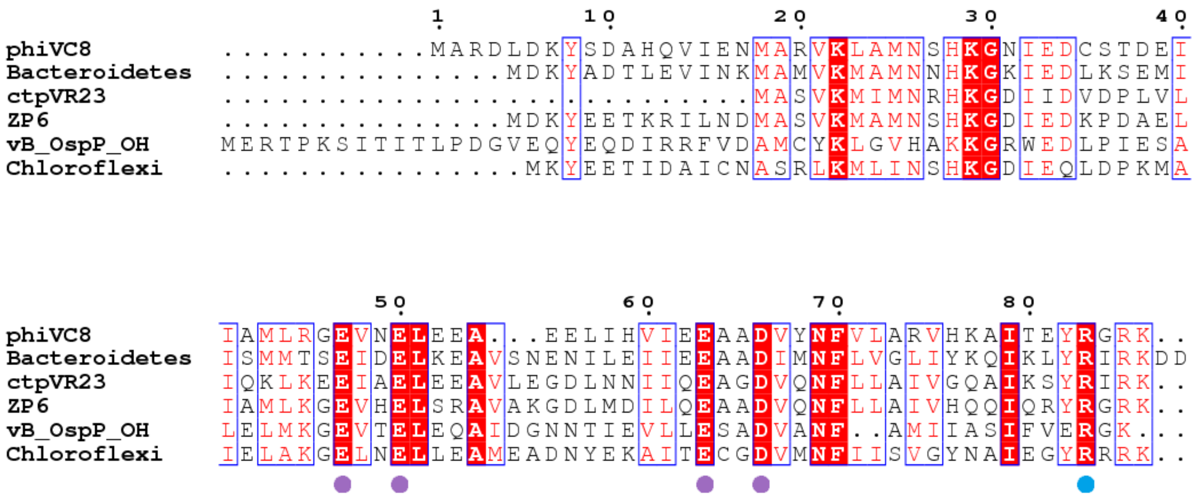


**Supplementary Figure S11. Sequence multialignment of MazZ-2 homologues (ϕVC8-like).** Numbering above the alignment refers to ϕVC8 MazZ. Putative catalytic residues are marked with lilac and blue dots, corresponding to S-2L MazZ E35, E38, E50, D53 and R83.

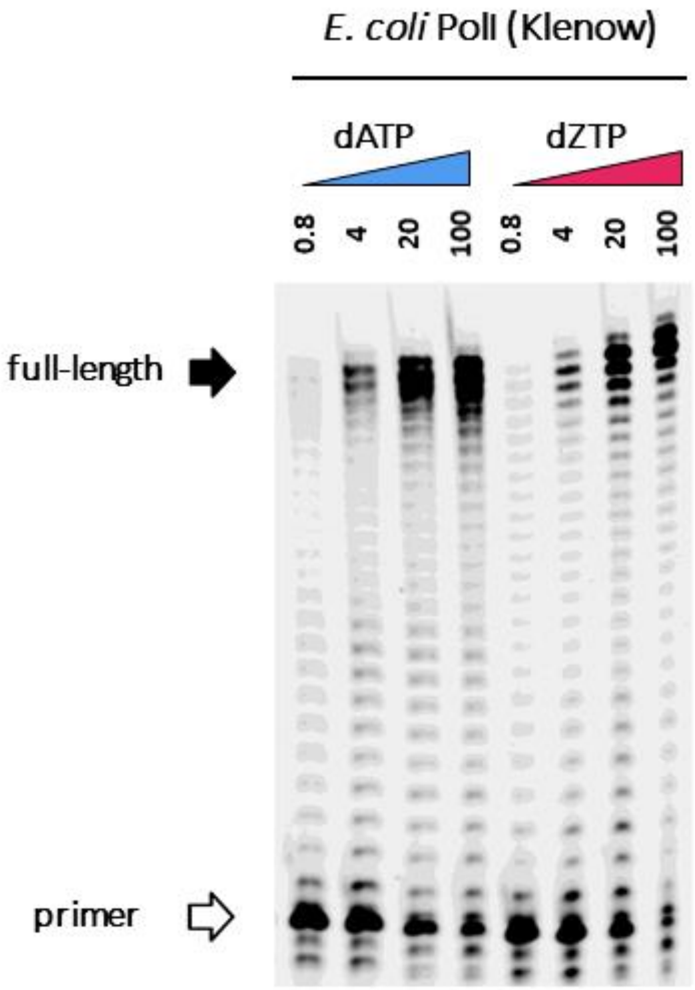


**Supplementary Figure S12. Results of DNA polymerase activity tests of *E. coli* Pol I (Klenow fragment) with either dATP (blue) or dZTP (magenta) as the incoming dNTP.** Nucleotide concentrations are given in µM concentration under the triangles.

| **Protein structure** | **PurZ + dGMP, dATP** | **MazZ + dGDP, $Mn^{2+}$, $SO_4^{2-}$** |
|---|---|---|
| PDB ID | **7ODX** | **7ODY** |
| *Cell parameters* | | |
| Space group | P $6_2$ 2 2 | P $2_1$ $2_1$ $2_1$ |
| *a*, *b*, *c* (Å) | 108.18, 108.18, 142.33 | 53.59, 91.37, 114.28 |
| *α*, *β*, *γ* (°) | 90.0, 90.0, 120.0 | 90.0, 90.0, 90.0 |
| Solvent content (%) | 58.8 | 56.1 |
| *Data statistics* | | |
| Resolution (Å) | 44.50 - 1.70 (1.74 - 1.70) | 48.52 - 1.43 (1.47 - 1.43) |
| Wavelength (Å) | 0.980130 | 1.127129 |
| Rmerge (%) | 8.1 (266.1) | 8.9 (193.8) |
| Completeness (%) | 99.8 (98.0) | 99.9 (98.3) |
| Multiplicity | 39.6 (39.4) | 13.1 (10.9) |
| $I/\sigma(I)$ | 34.1 (1.7) | 14.1 (1.1) |
| $CC_{1/2}$ | 1.000 (0.694) | 0.999 (0.579) |
| *Refinement* | | |
| Resolution (Å) | 42.33 - 1.70 | 48.45 - 1.43 |
| Unique reflections | 54,792 | 104,232 |
| $R_{work}/R_{free}$ (%) | 15.94/17.60 | 16.14/16.56 |
| *No. of non-hydrogen atoms* | | |
| Protein | 2702 | 2937 |
| Ligand | 53 | 168 |
| Ions | 0 | 54 |
| Water | 394 | 488 |
| Hydrogen atoms | No | Yes |
| *Protein geometry* | | |
| RMSD - bond lengths (Å) | 0.010 | 0.008 |
| RMSD - bond angles (°) | 1.00 | 0.88 |
| Ramachandran favored/outliers (%) | 96.84/0.00 | 98.92/0.00 |
| Rotamers favored/poor (%) | 98.24/0.00 | 97.00/0.00 |
| Clashscore | 2.57 | 3.23 |
| *B-factors ($Å^2$)* | | |
| Type | Anisotropic | Anisotropic |
| Protein | 30.71 | 25.55 |
| Ligand | 28.27 | 19.21 |

| | | |
|---|---|---|
| Ions | - | 38.74 |
| Water | 43.73 | 41.38 |

**Supplementary Table S1.** Diffraction data collection and Model Refinement statistics. Numbers in parenthesis refer to the highest-resolution shell.

| Protein name | Function | Start | End |
|---|---|---|---|
| DatZ | dATP triphosphohydrolase | 13934 | 13407 |
| MazZ | (d)GTP diphosphohydrolase | 14248 | 13931 |
| PurZ | N6-succino 2-amino deoxyadenylate synthase | 15308 | 14229 |
| Exonuclease VII | exonuclease | 17970 | 17011 |
| MarR | transcription repressor | 18735 | 18382 |
| Helicase SF2 | helicase | 18872 | 20137 |
| VRR nuclease | resolvase | 20147 | 20656 |
| PrimPol | DNA polymerase | 20825 | 23038 |

**Supplementary Table S2.** Position of replication-related protein genes on the new, S-2L genome sequence with high-coverage (MW334946). Genes *datZ*, *mazZ* and *purZ* are encoded in a highly compact way, overlapping at their very ends.

| Gene | Nucleotide sequence |
| --- | --- |
| *datZ* | ATGACACTCCAGATCACCGAGACCTACGAGCGCCTGAGGGCGTCCCACATCAGCCGGTGGGGATCGTCCAGACGACCTACCCGCAGAA<br>CATCGCCGAACACATGTGGCGCGTTTGGCTCCTGTGCCGGGACTGGGGCGCTGCCGCCGGCATGCCCCAGCACACGGTCCGCCAGGCCT<br>GCGAGTTTGCCCTGGTCCACGACCTGGCCGAGATCCGGACGGGCGACGCCCCGACGCCCCACAAGACCCCGGAGCTCAAGGAGCTCCTG<br>GCCGGCATCGAGGCCCAGATCGTCCCGGAGGTGGCCGAGCTCGAGGCGACCATGGCCCCGGAGGCCAGAGAGCTTTGGAAGTTCTGCGA<br>CACCGCCGAGGCCGTCCTGTTCCTCAAGGTCAACGGCCTGGGCGCCCACGCCTACGACGTCCAGCACCTGCTGATGGAGCAGATGAAAC<br>GGCGCCTGATGGACTCGGTGTTGGATGTGGAGGTGCAGGACGAGCTCATGTTCCAGTTCGAGCGGACGATCAAGAAGACGTGA |
| *mazZ* | ATGCCCGCTACCGTCGCTGAGCTCCAGGCGGAGATCGCCGCCTGGATCCACCCCCTGAACCCCGACCGCCGCCCCGGCGGCACCATCGC<br>CAAGCTCCTGGAGGAGATCGGGGAGTTGATCGCCAGCGACCGGGCCCACGACCCGCTCGAGGTGGCCGACGTCCTGATCCTGGCCCTCG<br>ACCTGGCGACGCTCCTGGGCGTCGACGTGACCGAGGCCATCCGCGCCAAGCTCGCCATCAACCGGGCCCGCTCCTGGCCCGAGCCGAT<br>AACGGCGCCATGCGCCACATCCCCGGTTCCGATACCCCCTCCTTCCCATGA |
| *purZ* | ATGCTGTCCATTCCCCCCTACTATCGCGTGAAGAACTGCAACCTGATCGTCGACTGCCAGTACGGCAGCACCGGCAAGGGGCTCCTGGC<br>CGGCTACCTGGGGGCGCTCGAGGCCCCGCAGGTGCTGTGCATGGCACCCAGCCCCAACGCCGGCCACACCCTGGTCGAGGAGGACGGCA<br>CCGCCCGCGTCCACAAGATGCTGCCCCTGGGCATCACCAGCCCCAGCCTTGAGCGGATCTACCTGGGCCCCGGCTCGGTGATCGACATG<br>GACCGGCTCCTAGAGGAGTACCTGGCCCTCCCCCGGCAGGTGGAGCTCTGGGTCCACCAGAACGCCGCCGTCGTCCTCCAGGAGCACCG<br>GGATGAGGAGGCCGCCGGGGGCCTGGCCCCAGGCTCGACCCGCAGCGGCGCCGGCTCGGCGTTTATCGCCAAGATCCGCCGCCGCCCTG<br>GGACGCTCCTGTTCGGTGAGGCCGTCCGGGATCACCCGCTCCACGGTGTCGTCCGGGTCGTCGACACCCGGACCGCCCAGGACATGCTG<br>TTTCGGACCCGGTCGATCCAGGCCGAGGGGTGCCAGGGCTACAGCCTGTCGGTCCACCACGGGGCCTACCCCTACTGCACCGCCCGGGA<br>CGTCACGACGGCCCAGCTGATCGCCGACTGCGGCCTGCCCTACGACGTCGCCCGGATCGCCCGGGTCGTCGGCTCGATGCGGACCTACC<br>CGATCCGGGTGGCCAACCGCCCGGAGGCCGGTGAGTGGAGCGGCCCCTGCTACCCCGACTCGGTCGAGTGCCAGTTCGCCGACCTGGGC<br>CTGGAGCAGGAGTACACCACCGTGACGAAGCTCCCCCGCCGGATCTTTACGTTTAGCGCCATCCAGGCGCACGAGGCCATCGCCCAGAA<br>CGGCGTGGACGAGGTGTTCCTCAACTTCGCCCAGTACCCGCCCAGCCTCGGGGCTCTCGAGGACATCCTCGACGCCATCGAGGCCAGGG<br>CGGAGGTGACCTACGTCGGCTTCGGCCCGAAGGTCACCGACGTCTACCACACCCCCACCCGGGCAGAGCTCGAAGGTTTGTATGCCCGC<br>TACCGTCGCTGA |
| *pplA* | ATGTCAACCCCCGCACCAGCCTTCGACCGGGACCAGATCCTCCTCCACCTGTCGCTCCTCCGGAAGGACATCGCCACGACCCGGTACCG<br>GGCGATCTGGCCCAGGCGAGAGGACAAGGTAAAAGCCTGGACGACGCCCCTGACCGGGGCCACGGTCCAGGACGCCGTCACCCAGGGAT<br>TCAACAGCTACATCGTCGTAGGCGACGGCGGCGACTCCGACGCCGAGATCACCAGTGTCAACGCCATCTTCGGCGAGTGGGACGACGGC<br>GACCTGGCCTGGCAGGTCGGCGCCTGGGAGGCCTGCGGCCTGCCGCGGCCGAGCTTCCAGCTGCGCACCGGGGGCAAGTCGATCCACCA<br>CTACTGGGTGTTCCACAGCCCTGTGGACGTCCCGGCCTGGACCGAGCTCCAGGCCCGCCTGATCGCCCTGGCCGGCTTCGACACGACGA<br>ACCGGAACCCCTCCCGGGTGATGCGCCTGGCCGGCTGCCCCCACCAGCGCACCGGGGAGGTGGCCCAGATCTTCAACGCGACCGGGGAG<br>CTCTACGACCCCGGGCAGATGCTGCAGGTCCTGCCCCCGGTGCCGATCGACCCGCCGGCTGCCGCCCCGGTGGCCCCGGGAGGTGCCCC<br>CAGTTCGATGGACGACATCCGGGCCGCCCTGGCCCAGATCCCACCCCGTCCCGGGGCAGGGAGCGGCACCTACGCCGAGTACCGCAACA<br>TCCTCTGGGGCCTGGTTAAGGCCGTCGAGGAGGCCGGCGGCACCCGGGACCAGGCCGTGGCCATGATGCAGGCGCACAGCCCCGAGGGC<br>TGGGATTGCGCCCAGGTGGCCCGCTCCGGGGGCAAGAAGATCAGCACCGGGACGTTCTGGTGGCATGCGATGTCCTACGGCTGGGCACC<br>GCCGAAGAAGGCCCCGGAGCCGCCGCCCCAGGCCCGCCAGGTGCCGGCCGTGGCCGCCGTGCTCCAGGCCGCAGAGGCCGCCCCTGGAA<br>CCGGCACCGAGCACGGCCCCTGGGCGCCGCTGCCCCCGGGCTGGCAGGGCACGAACAAGGAGGGCCTGCCCAGGGCCTCGCAGATCACC<br>ACCTACGAACTGGCCCTGCTGATGCAGGTCTCCCTGCGGGGGGTGCTCTGGCACAACGAGATGTCAGGCGAAGTCATGCACGGCAAGAC<br>GGCCCTCTCGCCGATCGAGCTCCAGATCGCCTACAGCCGGCTCGAGGGCCTCGGCTACAAGGTCACCAAGGAGAACGCCAAGACCGCCA<br>TCCTGCAGGCGTCGATCGCCGACCTGCGGCACCCCGTCCGGGAGTACCTCAACACCTGCACGACGCCCCTGCCCGACGAGGTCTGGGCC<br>GACATCGCCAACGCCCTGCTGGGCCCCGGGCACAGCGCGTTCGACTCCAGCGCCATCCGCAAGTGGCTGATCTTCGCCGTGGCCCGGGT<br>CTTCCAGCCCGGCTGCCCCTTCGGCTTCATGCTGGTGCTGGCTGGCGCCCAGCAGATGCACAAGACCCGGTTCTTTAACACCCTGGCCT<br>CAGACGAGTGGTTCCTGGGCGGATTCCAGCGGGGCCGCTCTGACACCGACGACCTGATTGCCCTGCACCGGTCCTGGATCACCGAGTGG<br>GGGGAGCTCGACGGCGGCCTCTCCAAGCACGACAGCGCCGAGCTCAAGGCGATGATCGACCGGAAGGTGGACGTGCTCCGGAGGCCCTA<br>CGCCGCCACGCACGAAAGCTGCCCCCGGAGCTTCGTCCTCTGCGGGACGACGAACCGCCGGGATGGGCTCTTCACCGACCCGACCGGCA<br>ACAGGCGGTACGTGGTCGTGCCCGTCAACCAGCGGATCGACAGCGAGCGCCTGGAGCAGATGCGAGACCAGATCTGGGCAACCGCCCTC<br>CGGGAGTACCGCAGCGGCAAGCTCTGGTACCTCGACGAGGAGGAGCTGGAGATCAACGCGAAACGCAACAAGGGCCTTGAGGTGGAGGA<br>CGCCTGGGTGGGGACGATCCAGATGCACCTGAATAGCTCGATCGACCTGGAGCGGCTGACCGACGGGCGCTACGGCATCAACATCGAGT<br>CAGTCTACCTCAAGATCGAGCCCGAGGTGGACGCCGTGGCCCGGGCTTCGGAAAGCGGATCCGGGACACCATGCTGAGCCTGGGCTGG<br>GAGCCCGTGCGGCTGCGTCTCGCCAGCGACCCGAGCGGCAACCCGGTGAGGCGTTGGGCGCCCGTCCAGGGGGGGTAG |

**Supplementary Table S3.** Nucleotide sequences of *datZ*, *mazZ*, *purZ* and *pplA* native genes (GenBank MW334946).

| Gene | Nucleotide sequence |
|---|---|
| *datZ* | ATGACACTGCAGATTACCGAAACCTATGAACGTCTGCGTGCAAGCCATATTAGCCGTTGGGGTATTGTTCAGACCACCTATCCGCAGAA<br>TATTGCAGAACATATGTGGCGTGTTTGGCTGCTGTGTCGTGATTGGGGTGCAGCAGCAGGTATGCCGCAGCATACAGTTCGTCAGGCAT<br>GTGAATTTGCACTGGTTCATGATCTGGCAGAAATTCGTACCGGTGATGCACCGACACCGCATAAAACACCGGAACTGAAAGAACTGCTG<br>GCAGGTATTGAAGCACAGATTGTTCCGGAAGTTGCAGAACTGGAAGCAACCATGGCACCGGAAGCACGTGAACTGTGGAAATTTTGTGA<br>TACCGCAGAAGCAGTTCTGTTCCTGAAAGTTAATGGTCTGGGTGCACATGCATATGATGTTCAGCATCTGCTGATGGAACAAATGAAAC<br>GTCGTCTGATGGATAGCGTTCTGGATGTTGAAGTTCAGGATGAACTGATGTTTCAGTTTGAACGCACCATCAAAAAGACCTAA |
| *mazZ* | ATGCCTGCAACCGTTGCCGAACTGCAGGCAGAAATTGCAGCCTGGATTCATCCGCTGAATCCGGATCGTCGTCCTGGTGGCACCATTGC<br>AAAACTGCTGGAAGAAATCGGTGAACTGATTGCAAGCGATCGTGCACATGATCCGCTGGAAGTTGCAGATGTTCTGATTCTGGCACTGG<br>ATCTGGCAACCCTGCTGGGTGTTGATGTTACCGAAGCAATTCGTGCCAAACTGGCAATTAATCGTGCACGTAGCTGGGCACGTGCAGAT<br>AATGGTGCAATGCGTCATATTCCGGGTAGCGATACCCCGAGCTTTCCGTAA |
| *purZ* | ATGCTGAGCATTCCGCCTTATTATCGTGTGAAAAATTGCAACCTGATTGTGGATTGTCAGTATGGTAGCACCGGTAAAGGTCTGCTGGC<br>AGGTTATCTGGGTGCACTGGAAGCACCGCAGGTTCTGTGTATGGCACCGAGTCCGAATGCAGGTCATACCCTGGTTGAAGAGGATGGCA<br>CCGCACGTGTTCATAAAATGCTGCCGCTGGGTATTACCAGTCCGAGCCTGGAACGTATTTATCTTGGTCCGGGTAGCGTTATTGATATG<br>GATCGTCTGCTGGAAGAATATCTGGCACTGCCTCGTCAGGTTGAACTGTGGGTTCATCAGAATGCAGCAGTTGTTCTGCAAGAACATCG<br>TGATGAAGAAGCAGCAGGCGGTCTGGCACCGGGTAGCACCCGTAGCGGTGCAGGTAGCGCATTTATTGCAAAAATTCGTCGTCGTCCGG<br>GTACACTGCTGTTTGGTGAAGCAGTTCGTGATCATCCGCTGCATGGTGTTGTTCGTGTTGTTGATACCCGTACCGCACAGGATATGCTG<br>TTTCGTACCCGTAGCATTCAGGCAGAAGGTTGTCAGGGTTATAGCCTGAGCGTTCATCATGGTGCATATCCGTATTGTACAGCACGTGA<br>TGTTACCACCGCACAGCTGATTGCAGATTGTGGTCTGCCGTATGATGTTGCACGTATTGCACGTGTTGTGGGTAGCATGCGTACCTATC<br>CGATTCGTGTTGCAAATCGTCCGGAAGCCGGTGAATGGTCAGGTCCGTGTTATCCGGATTCAGTTGAATGTCAGTTTGCAGATCTGGGC<br>CTTGAACAAGAATATACCACCGTTACCAAACTGCCACGTCGCATTTTTACCTTTAGCGCAATTCAGGCACATGAAGCAATTGCACAGAA<br>TGGTGTTGATGAAGTGTTTCTGAATTTTGCACAGTATCCGCCTAGCCTGGGAGCCCTGGAAGATATTCTGGATGCAATTGAAGCACGTG<br>CCGAAGTTACCTATGTTGGTTTTGGTCCGAAAGTTACCGATGTGTATCATACCCCGACACGTGCAGAACTGGAAGGTCTGTATGCACGT<br>TATCGTCGTTAA |
| *pplA* | ATGAGCACACCGGCACCGGCATTTGATCGTGATCAGATTCTGCTGCATCTGAGCCTGCTGCGTAAAGATATTGCAACCACACGTTATCG<br>TGCAATTTGGCCTCGTCGTGAAGATAAAGTTAAAGCATGGACCACACCGCTGACCGGTGCAACCGTTCAGGATGCAGTTACCCAGGGTT<br>TTAATAGCTATATCGTTGTTGGTGATGGTGGTGATAGTGATGCAGAAATTACCAGCGTTAATGCCATTTTTGGTGAATGGGATGATGGT<br>GATCTGGCATGGCAGGTTGGTGCATGGGAAGCATGTGGTCTGCCTCGTCCGAGCTTTCAGCTGCGTACCGGTGGTAAAAGCATTCATCA<br>TTATTGGGTTTTTCACAGTCCGGTTGATGTTCCGGCATGGACCGAACTGCAGGCACGTCTGATTGCACTGGCAGGTTTTGATACCACCA<br>ATCGTAATCCGAGCCGTGTTATGCGTCTGGCAGGCTGTCCGCATCAGCGCACCGGTGAAGTTGCACAGATTTTCAATGCAACCGGTGAA<br>CTGTATGATCCGGGTCAGATGCTGCAGGTTCTGCCTCCGGTTCCGATTGATCCGCCTGCAGCAGCTCCGGTTGCGCCTGGTGGTGCACC<br>GAGCAGCATGGATGATATTCGTGCAGCACTGGCACAGATTCCGCCTCGTCCTGGTGCAGGTAGCGGCACCTATGCAGAATATCGTAATA<br>TTCTGTGGGGTTTAGTTAAAGCCGTTGAAGAGGCAGGCGGTACACGTGATCAGGCAGTTGCAATGATGCAGGCACATAGTCCGGAAGGT<br>TGGGATTGTGCACAGGTTGCACGTAGTGGTGGCAAAAAAATCAGCACCGGTACATTTTGGTGGCATGCAATGAGCTATGGTTGGGCACC<br>GCCTAAAAAAGCACCGGAACCGCCTCCGCAGGCACGCCAGGTTCCAGCAGTTGCAGCAGTTCTGCAGGCAGCAGAAGCAGCCCCTGGTA<br>CAGGCACCGAACATGGTCCGTGGGCTCCGCTGCCTCCTGGTTGGCAGGGCACCAATAAAGAAGGTCTGCCACGCGCAAGCCAGATTACC<br>ACCTATGAACTGGCACTGCTGATGCAGGTTAGCCTGCGTGGTGTTCTGTGGCATAATGAAATGAGCGGTGAAGTAATGCATGGTAAAAC<br>CGCACTGAGCCCGATTGAACTGCAGATTGCATATAGCCGTCTGGAAGGTCTGGGTTATAAAGTGACCAAAGAAAATGCAAAAACCGCAA<br>TCCTGCAGGCAAGCATTGCCGATCTGCGTCATCCGGTTCGTGAATATCTGAATACCTGTACAACCCCTCTGCCGGATGAAGTTTGGGCA<br>GATATTGCCAATGCACTGTTAGGTCCGGGTCATAGCGCATTTGATAGCAGCGCAATTCGTAAATGGCTGATCTTTGCAGTTGCACGTGT<br>TTTTCAGCCTGGTTGTCCGTTTGGTTTTATGCTGGTGCTGGCAGGCGCACAGCAGATGCATAAAACACGCTTTTTTAACACCCTGGCAT<br>CCGATGAATGGTTTTTAGGTGGTTTTCAGCGTGGTCGTAGCGATACCGATGATCTGATTGCCCTGCATCGTAGCTGGATTACCGAATGG<br>GGTGAACTGGATGGTGGTCTGAGCAAACATGATAGCGCAGAACTGAAAGCAATGATTGATCGTAAAGTTGATGTGCTGCGTCGTCCGTA<br>TGCAGCAACCCATGAAAGCTGTCCGCGTAGCTTTGTTCTGTGTGGTACAACCAATCGTCGTGATGGTCTGTTTACCGATCCGACCGGTA<br>ATCGTCGTTATGTTGTTGTTCCGGTTAATCAGCGTATTGATAGCGAACGTCTGGAACAAATGCGCGATCAGATTTGGGCCACCGCACTG<br>CGCGAATATCGTTCAGGTAAACTGTGGTATCTGGATGAAGAGGAACTGGAAATTAATGCCAAACGCAATAAAGGTCTGGAAGTTGAAGA<br>TGCATGGGTTGGCACCATTCAGATGCACCTGAACAGCAGCATTGATCTGGAACGTCTGACCGATGGTCGTTATGGTATTAACATTGAAA<br>GCGTGTACCTGAAAATTGAACCGGAAGTTGGTCGTCGTGGTCCTGGTTTTGGTAAACGTATTCGTGATACCATGCTGAGCTTAGGTTGG<br>GAACCTGTTCGTCTGCGCCTGGCAAGCGATCCGAGTGGTAATCCGGTGCGTCGCTGGGCACCTGTTCAAGGTGGTTAA |

**Supplementary Table S4.** Nucleotide sequences of *datZ*, *mazZ*, *purZ* and *pplA* codon-optimized genes.

| Protein | Protein sequence |
|---|---|
| DatZ | MTLQITETYERLRASHISRWGIVQTTYPQNIAEHMWRVWLLCRDWGAAAGMPQHTVRQACEFALVHDLAEIRTGDAPTPHKTPELK<br>ELLAGIEAQIVPEVAELEATMAPEARELWKFCDTAEAVLFLKVNGLGAHAYDVQHLLMEQMKRRLMDSVLDVEVQDELMFQFERTI<br>KKT- |
| MazZ | MPATVAELQAEIAAWIHPLNPDRRPGGTIAKLLEEIGELIASDRAHDPLEVADVLILALDLATLLGVDVTEAIRAKLAINRARSWA<br>RADNGAMRHIPGSDTPSFP- |
| PurZ | MLSIPPYYRVKNCNLIVDCQYGSTGKGLLAGYLGALEAPQVLCMAPSPNAGHTLVEEDGTARVHKMLPLGITSPSLERIYLGPGSV<br>IDMDRLLEEYLALPRQVELWVHQNAAVVLQEHRDEEAAGGLAPGSTRSGAGSAFIAKIRRRPGTLLFGEAVRDHPLHGVVRVVDTR<br>TAQDMLFRTRSIQAEGCQGYSLSVHHGAYPYCTARDVTTAQLIADCGLPYDVARIARVVGSMRTYPIRVANRPEAGEWSGPCYPDS<br>VECQFADLGLEQEYTTVTKLPRRIFTFSAIQAHEAIAQNGVDEVFLNFAQYPPSLGALEDILDAIEARAEVTYVGFGPKVTDVYHT<br>PTRAELEGLYARYRR- |
| PrimPol | MSTPAPAFDRDQILLHLSLLRKDIATTRYRAIWPRREDKVKAWTTPLTGATVQDAVTQGFNSYIVVGDGGDSDAEITSVNAIFGEW<br>DDGDLAWQVGAWEACGLPRPSFQLRTGGKSIHHYWVFHSPVDVPAWTELQARLIALAGFDTTNRNPSRVMRLAGCPHQRTGEVAQI<br>FNATGELYDPGQMLQVLPPVPIDPPAAAPVAPGGAPSSMDDIRAALAQIPPRPGAGSGTYAEYRNILWGLVKAVEEAGGTRDQAVA<br>MMQAHSPEGWDCAQVARSGGKKISTGTFWWHAMSYGWAPPKKAPEPPPQARQVPAVAAVLQAAEAAPGTGTEHGPWAPLPPGWQGT<br>NKEGLPRASQITTYELALLMQVSLRGVLWHNEMSGEVMHGKTALSPIELQIAYSRLEGLGYKVTKENAKTAILQASIADLRHPVRE<br>YLNTCTTPLPDEVWADIANALLGPGHSAFDSSAIRKWLIFAVARVFQPGCPFGFMLVLAGAQQMHKTRFFNTLASDEWFLGGFQRG<br>RSDTDDLIALHRSWITEWGELDGGLSKHDSAELKAMIDRKVDVLRRPYAATHESCPRSFVLCGTTNRRDGLFTDPTGNRRYVVVPV<br>NQRIDSERLEQMRDQIWATALREYRSGKLWYLDEEELEINAKRNKGLEVEDAWVGTIQMHLNSSIDLERLTDGRYGINIESVYLKI<br>EPEVGRRGPGFGKRIRDTMLSLGWEPVRLRLASDPSGNPVRRWAPVQGG- |

**Supplementary Table S5.** Protein sequences of DatZ, MazZ, PurZ and PrimPol.
